# mRNA/LNP-mediated expression of a ligand-independent Tie2 pathway activator, COMP-Tie2*, enables sustained endothelial stabilization

**DOI:** 10.64898/2026.11.10.765541

**Authors:** Katrin Radloff, Charlotte Dunne, Oliver Keil, Ansgar Santel, Jörg Kaufmann, Daniel Tondera

## Abstract

Activation of the angiopoietin/Tie2 signaling axis is a validated strategy for endothelial stabilization. However, therapeutic Tie2 activation via angiopoietin-1 (Ang1) or Ang1-derived agonists is intrinsically limited due to several mechanism including ligand instability, competitive Ang2 inhibition, Tie2 ectodomain cleavage, VE-PTP-mediated Tie2 dephosphorylation, and ligand-induced receptor degradation. All these inhibitory mechanisms restrict both magnitude as well as duration of therapeutic Tie2 signaling, particularly under inflammatory conditions such as acute respiratory distress syndrome (ARDS) or sepsis. Here, we describe a novel ligand-independent strategy for sustained Tie2 activation based on engineered membrane-anchored Tie2 variants. By combining intracellular Tie2 kinase domains harboring naturally occurring autoactivating mutations with heterologous membrane-targeting domains, we generated truncated, constitutively active Tie2 variants that bypass extracellular ligand regulation entirely. mRNA-mediated expression of these constructs induced robust and persistent Tie2 autophosphorylation and downstream Akt–FOXO1 signaling in pulmonary endothelial cells, which exceeded COMP-Ang1–mediated activation in both strength and duration. Systemic administration of COMP-Tie2* mRNA using lung-targeted lipid nanoparticles resulted in selective pulmonary Tie2 receptor engagement *in vivo.* Collectively, these data establish LNP-mediated mRNA delivery of engineered recombinant receptors for intrinsic Tie2 activation as a promising novel approach for endothelial stabilization under inflammatory conditions.

**Graphical abstract:** 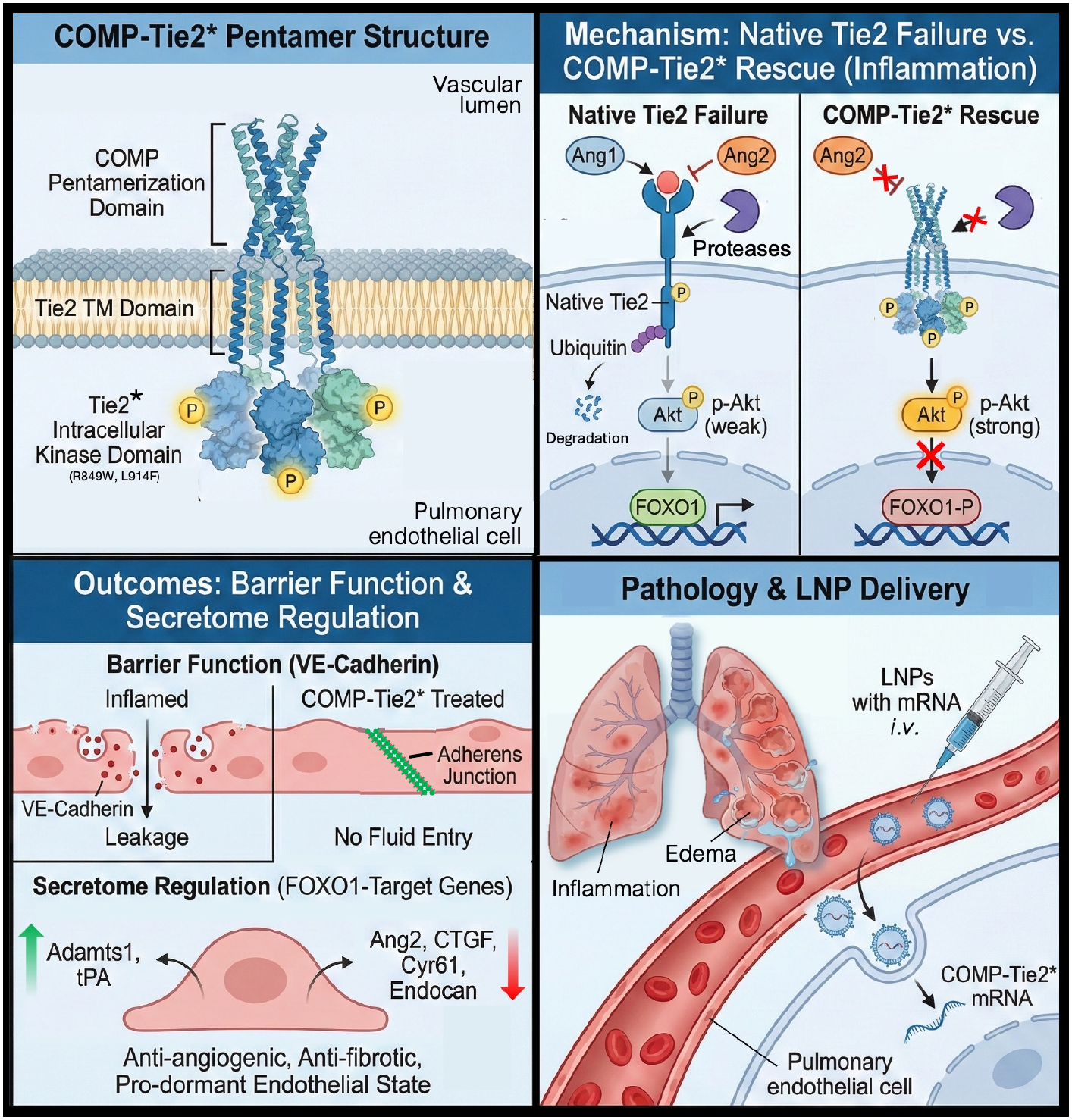

## Introduction

Loss of endothelial integrity in organs with continuous microvascular architecture such as the lung increases vascular permeability, causing pulmonary edema, disrupting microvascular perfusion, and ultimately leading to organ failure, with consequent morbidity and mortality^1,2^. Endothelial barrier dysfunction is a defining pathological feature of several inflammatory vascular diseases, including acute respiratory distress syndrome (ARDS), or sepsis. Restoration of pulmonary endothelial barrier function has therefore emerged as a central therapeutic objective in inflammatory disease^3^.

The angiopoietin/Tie2 signaling axis (pathway?) is one of the most important endogenous systems governing endothelial stability^4–9^. Under homeostatic conditions, angiopoietin-1 (Ang1, ANGPT1) activates the receptor tyrosine kinase Tie2 (TEK), promoting endothelial quiescence, which in turn leads to stabilization of intercellular junctions, suppression of inflammatory gene expression, and maintenance of vascular integrity^10–14^. In contrast, inflammatory conditions are characterized by elevated angiopoietin-2 (Ang2, ANGPT2) released from endothelial Weibel-Palade bodies. Ang2 functions as a context-dependent antagonist of Tie2 signaling and drives both endothelial activation and vascular leakage^15,16^. Clinical studies have consistently demonstrated that increased circulating Ang2 levels strongly correlate with ARDS disease severity and mortality, underscoring the central role of Tie2 signaling loss in disease pathophysiology^17–20^.

Native Ang1 is poorly soluble, highly aggregation-prone, and difficult to produce as a recombinant protein in a stable and scalable manner^21^. Considerable efforts have been centered on therapeutic activation of Tie2 using recombinant Ang1-derived agonists^21–25^, However, the clinical translation of ligand-based Tie2 therapies has proven challenging. Even engineered Ang1 derivatives such as COMP-Ang1 remain complex multimeric proteins with limited manufacturability and formulation stability, while improving controlled oligomerization and bioactivity. These challenges have significantly constrained the development of recombinant Ang1-based biologics for clinical use and have led to the development of alternative Tie2 activators such as agonistic Tie2 antibodies^26,27^, Tie2 binding peptides^28,29^ or Ang2-inhibiting antibodies^30–32^.

Beyond production-related limitations, several intrinsic biological mechanisms further restrict the efficacy of ligand-dependent Tie2 activation *in vivo*. During inflammation, protein levels of Ang2 are elevated and compete with Ang1 for Tie2 binding while actively suppressing receptor signaling^33,34^. In parallel, inflammatory stimuli promote proteolytic cleavage of the Tie2 ectodomain, resulting in receptor shedding and generation of soluble Tie2 fragments which further sequester circulating Ang1^35–38^. This reduction in functional surface Tie2 profoundly limits the responsiveness of the endothelium to ligand-based agonists specifically under conditions in which Tie2 activation would be most desirable. In addition, Tie2 signaling is tightly regulated at the receptor level via vascular endothelial protein tyrosine phosphatase (VE-PTP), which dephosphorylates Tie2 and limits signaling duration^39,40^. Pharmacological inhibition of VE-PTP was therefore proposed as an alternative strategy to sustain Tie2 activation^41^.

However, despite promising preclinical data, VE-PTP inhibitors have not met therapeutic expectations^42^, likely due to incomplete pathway engagement, context-dependent effects, and limited efficacy in severe inflammatory settings. Taken together, these findings indicate that Tie2 signaling is governed by multiple, overlapping regulatory mechanisms that are not easily overcome by ligand augmentation or phosphatase inhibition alone.

In addition, even when Tie2 activation is achieved, ligand-mediated receptor signaling remains inherently transient. Ang1 stimulation, as is typical for receptor tyrosine kinases, inducesTie2 ubiquitination and internalization followed by receptor degradation, which progressively depletes total Tie2 protein and attenuates signaling over time^43–46^. Thus, ligand-based Tie2 activation is not only antagonized extracellularly but is also self-limiting at the receptor level.

In our previous work, we demonstrated that lung-targeted cationic lipid nanoparticle (cLNP) mediated delivery of COMP-Ang1 mRNA to the pulmonary endothelium enables localized pulmonary Tie2 activation and endothelial stabilization *in vivo*^44^. This approach overcame several limitations of recombinant protein delivery and demonstrated that mRNA-based Tie2 modulation is a feasible therapeutic strategy in the lung vasculature. However, these studies also revealed rapid attenuation of Tie2 signaling consistent with transient expression and rapid clearance of the COMP-Ang-1 together with a rapid ligand-driven receptor turnover, underscoring the fundamental biological constraints of therapeutic Ang1-dependent receptor activation.

Together, these observations suggest that durable restoration of Tie2 signaling cannot be achieved solely by increasing ligand availability or enhancing ligand potency. Instead, sustained endothelial stabilization may require a fundamentally different approach that bypasses extracellular ligand regulation and directly engages receptor-intrinsic signaling mechanisms.

We therefore hypothesized that stable Tie2 pathway activation might be achieved via mRNA-mediated expression of Ang1/Ang2-independent Tie2 receptor variants. We aimed at overcoming the combined limitations of ligand producibility, receptor shedding and competitive antagonism by engineering constitutively active, membrane-anchored constitutively active Tie2 constructs which no longer rely on ligand binding. In the present study, we describe the molecular design and functional characterization of such Tie2 activators and evaluate their capacity to induce sustained pulmonary microvascular endothelial Tie2 signaling in vitro and in vivo.

## Results

### Evaluation of synthetic Tie2 mRNAs for ligand-independent Tie2 receptor signaling

On order to identify new molecular tools to allow for an Ang1/Ang2-independent activation of Tie2 signaling, we first focused on naturally occurring autosomal dominant hyperactivating Tie2 mutations that have been described in somatic and germline vascular disorders. Among these, the substitutions R849W and L914F within the intracellular kinase domain have been reported to confer constitutive Tie2 activity^47–50^. Based on this information, we generated PTXmRNA constructs encoding wild-type Tie2, the individual point mutants Tie2^R849W^ and Tie2^L914F^, as well as a combined double mutant (Tie2*), in which both substitutions were present (Figure 1A). To assess ligand dependency, the respective Tie2 mRNAs were transfected into Tie2-negative HeLa cells for 24 h, followed by treatment with or without recombinant human Ang1 for 15 min.

**Figure 1.**
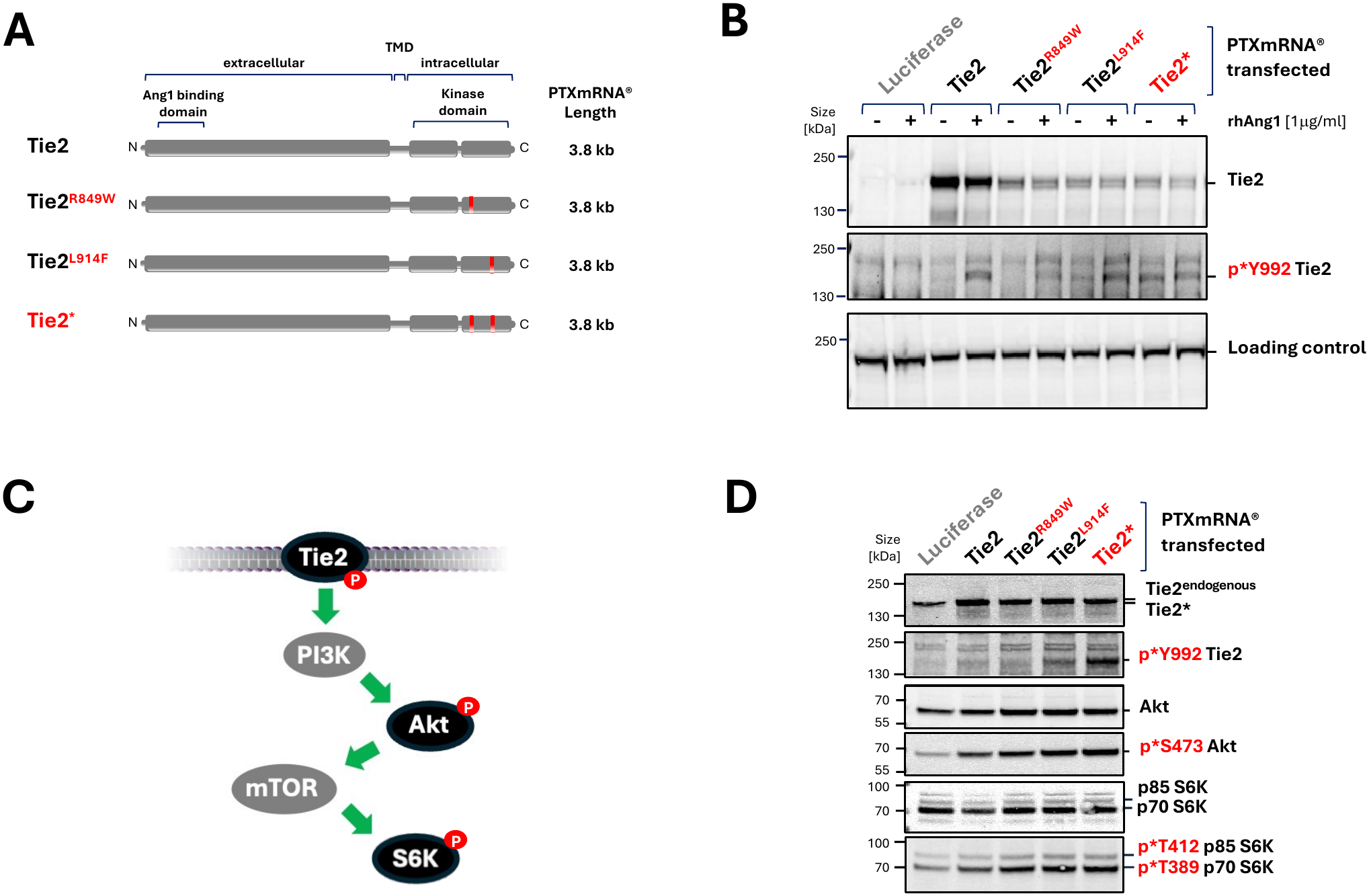
Identification and engineering of ligand-independent Tie2 receptor variants. **(A)** Schematic representation of the protein domain structure of human Tie2 and naturally occurring Tie2 variants carrying the R849W or L914F substitutions, as well as the combined double mutant Tie2*. The Ang1-binding extracellular domain, transmembrane domain (TMD), and intracellular kinase domain are indicated. **(B)** Assessment of Ang1 dependency of Tie2 variants in Tie2-negative HeLa cells. Cells were transfected with PTXmRNA® encoding wild-type Tie2 or indicated Tie2 variants and after 24h cells were stimulated for 15 min with recombinant human Angiopoietin-1. Tie2 activation was analyzed by Western blotting using phospho-specific antibodies recognizing Tie2 Y992. **(C)** Simplified canonical Tie2-Akt-S6K pathway and its components used to analyze Tie2 pathway activation (black). **(D)** Expression and downstream signaling of Tie2 variants in Tie2-positive human pulmonary microvascular endothelial cells (HPMECs). Serum-starved cells were transfected with mRNAs encoding wild-type Tie2 or Tie2 variants for 6h and analyzed for Tie2 activation (p*Y992) and downstream Akt phosphorylation (p*S473) and p70 and p85 S6 Kinase phosphorylation (p*T389/p*T412).

Lysates from cells transfected with mRNA encoding luciferase were used as negative control. All Tie2 constructs were expressed at detectable levels whereas all three autoactivating variants consistently showed reduced steady-state protein abundance compared with wild-type Tie2 (Figure 1B). The reduced expression of the autoactivating Tie2 mutations suggest an accelerated activation-dependent receptor turnover as described elsewhere^43,44^. While wild-type Tie2 and both single mutants remained responsive to Ang1 stimulation, the Tie2* double mutant exhibited high basal phosphorylation at Y992 which was not further increased by Ang1 treatment (Figure 1B, middle panel). These data identify Tie2* as a fully Ang1-independent Tie2 pathway activator. We next evaluated Tie2 signal transduction in a physiologically more relevant endothelial cell context by analyzing activation of the canonical Tie2-Akt-S6 Kinase pathway (Figure 1C). Serum-starved Tie2-positive human pulmonary microvascular endothelial cells (HPMECs) were transfected with the respective Tie2 mRNAs and analyzed for receptor activation and downstream signaling. Tie2* induced the strongest basal phosphorylation of Tie2 at Y992, confirming robust autoactivation in endothelial cells. However, despite variations in receptor phosphorylation intensity, all activating variants induced a similarly elevated downstream PI3K-Akt-mTOR-S6K signaling response, suggesting that maximal Akt and S6K activation can occur across a broad range of Tie2 phosphorylation levels (Figure 1D).

### Engineering of truncated membrane-targeted Tie2* variants for more potent Tie2 pathway activation

We next explored whether the extracellular Tie2 domain could be removed to generate shorter, ligand-independent constructs, since Tie2* activation was independent of Ang1 binding. Two truncated Tie2* variants were designed and engineered (Figure 2A). In the first construct (myr-Tie2*), the ectodomain and transmembrane domain of Tie2* were replaced by the N-terminal myristylation motif of Src kinase, enabling plasma membrane insertion following post-translational myristylation. In the second construct (COMP-Tie2*), a MCP-1–derived signal peptide and the pentamerization domain of cartilage oligomeric matrix protein (COMP) were fused upstream of the Tie2* transmembrane and kinase domains, thus enabling membrane localization combined with extracellular pentameric clustering. To assess the potential of myr-Tie2* and COMP-Tie2* for activating Tie2 pathway in the absence of ligands, we analyzed the canonical Akt-FOXO1 signaling induced by these truncated Tie2* variants (Figure 2B) and the phosphorylation status of additional Tie2 phosphorylation sites within the truncated Tie2* variants (Figure 2C). Serum-starved HPMECs were transfected with increasing amounts of myr-Tie2* or COMP-Tie2* mRNA, and lysates were analyzed after 8 hours. A strong, dose-dependent expression of both variants was detected using a C-terminal Tie2 antibody (Figure 2D, Panel 2). In addition to Y992 (Figure 2D, Panel 3), a strongly dose-dependent phosphorylation was also observed for both constructs at equal levels at Tie2 residues Y1102 and S1119 (Figure 2D, Panel 4 and 5). Both respective phosphorylation-sites are characteristically very difficult to detect on endogenous Tie2 by Western blotting (Figure 2B).

**Figure 2.**
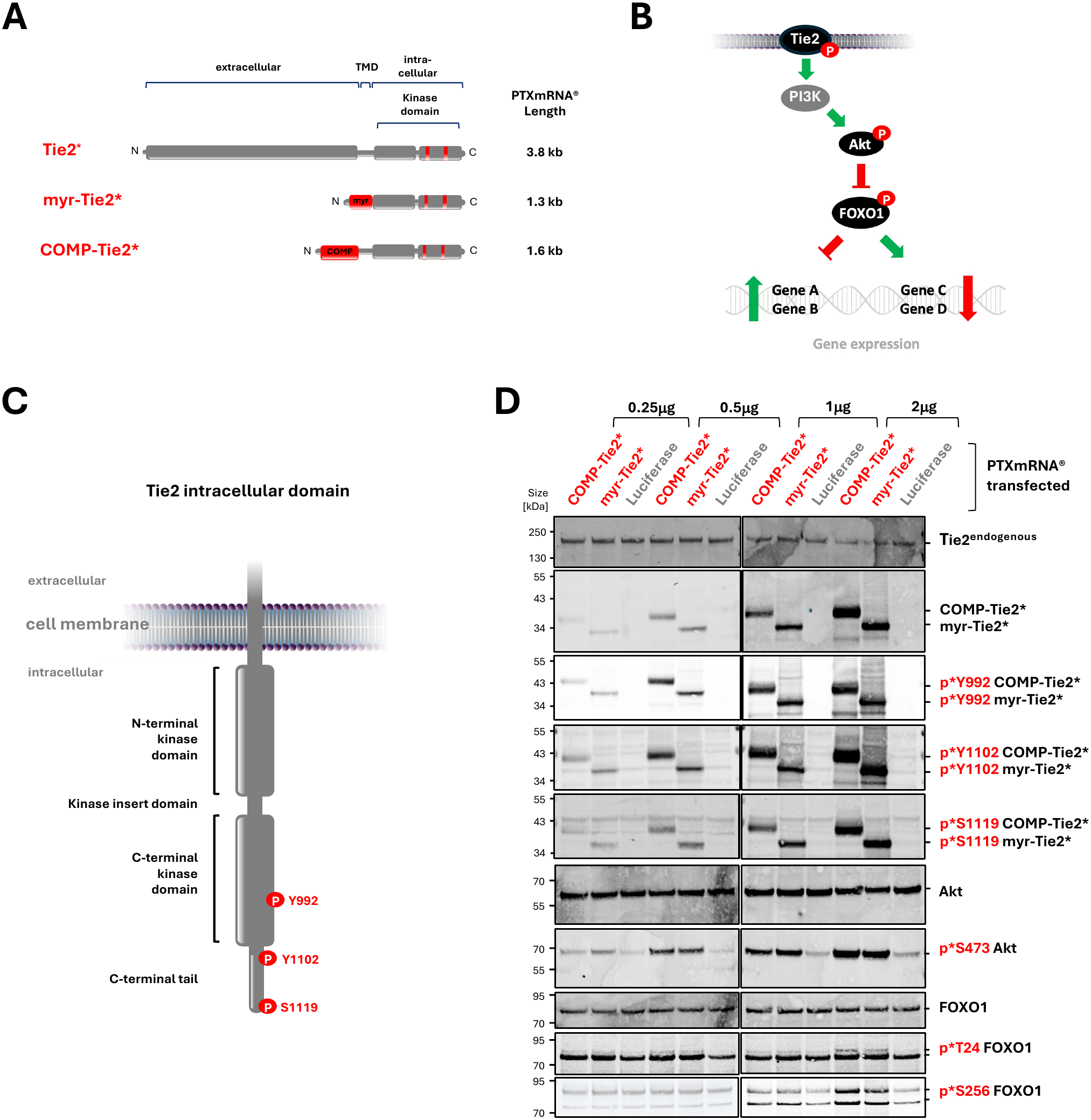
Identification and engineering of ligand-independent, short Tie2 receptor variants. **(A)** Schematic representation of engineered truncated Tie2 fusion constructs. The Tie2 extracellular domain and transmembrane domain was replaced by either an N-terminal myristylation motif from Src kinase (myr-Tie2*) or the Tie2 extracellular domain was replaced by an N-terminal MCP-1-derived secretion signal and a pentamerization domain derived from cartilage oligomeric matrix protein (COMP, TSP5) (COMP-Tie2*). **(B)** Canonical Tie2-Akt-FOXO1 pathway components used for signaling analysis and **(C)** schematic Tie2 ICD structure and phosphorylation sites used to analyze Tie2 pathway activation. **(D)** Dose-dependent expression and phosphorylation of engineered Tie2 variants and downstream pathway activation in HPMECs. Serum-starved cells were transfected with increasing amounts of PTXmRNA® encoding myr-Tie2* or COMP-Tie2* using Messenger Max. Western blot analysis from protein lysates was performed after 8h to assess Tie2 autophosphorylation at Y992, Y1102, and S1119, Akt activation at S473, FOXO1 inhibition at T24 and S256. Detection of endogenous Tie2, Akt, and FOXO1 served as loading controls.

This dose-dependent increase in Tie2 autophosphorylation correlated with a corresponding dose-dependent activation of Akt, as evidenced by Akt phosphorylation at S473. (Figure 2D, Panel 7). A similar correlation was observed for FOXO1 phosphorylation at T24 and S256 (Figure 2D, Panel 9 and 10), consistent with functional inhibition of this transcription factor upon Tie2 activation.

### Time dependent comparison of Tie2 signaling induced by COMP-Ang-1, Tie2* with truncated membrane-targeted Tie2* variants in HPMECs

To compare signaling potency, serum-starved HPMECs were transfected with mRNAs encoding full-length Tie2*, myr-Tie2*, COMP-Tie2*, or COMP-Ang1 as a ligand-based comparator. COMP-Ang1 expression and secretion into the supernatant was detected using an Ang1-specific antibody (Figure 3A, Supplementary Figure S1), while endogenous Tie2 and Tie2* were detected using N-terminal Tie2 antibodies unable to detect N-terminal truncated myr-Tie2* and COMP-Tie2* (Figure 3B, Panel 1 and 2). The N-terminally truncated Tie2* variants were detected using p*Y922 antibodies or C-terminal Tie2 antibodies (Figure 3C, Panel 1 and 2). However, due to their limited sensitivity, these antibodies failed to detect endogenous Tie2 or Tie2*. All constructs were detectably expressed within 2 hours post-transfection, although full-length Tie2*, representing the longest mRNA, showed the lowest early expression levels and peaked at 6 hours (Figure 3B, Lane 16). COMP-Ang1 expression and secretion increased over time (Figure 3A, Supplementary Figure S1), and induced rapid, but low Tie2 phosphorylation which decreased slightly over time together with reduced levels of endogenous Tie2 (Figure 3B, Panel 3). Tie2* expression was low at the beginning but increased over time (Figure 3B, Panel 1 and 2) to a level higher than what was initially achieved by COMP-Ang1, analogous to levels of phosphorylated Tie2* (Figure 3B, Panel 3). In contrast, both truncated Tie2* variants were highly expressed and induced robust Tie2 Y992 phosphorylation within 2 hours (Figure 3C, Panel 3), greatly surpassing the level of activation of endogenous Tie2 achieved by COMP-Ang1 or Tie2*. The level of Tie2 activation was directly reflected by the level of Akt activation indicated by S473 phosphorylation (Figure 3D, Panel 2). Direct comparison of the two short Tie2* variants with COMP-Ang1 and full-length Tie2* reveals that both truncated constructs induce markedly stronger Akt activation than ligand-mediated Tie2 stimulation or full length Tie2* variants. Surprisingly, analysis of endogenous Tie2 using antibodies recognizing different Tie2 epitopes (Figure 3, right) showed that myr-Tie2*, in contrast to COMP-Tie2*, is associated with a reduction in endogenous Tie2 levels (Figure 3B, Panel 1, Lane 17 and 23) to a level similar to the activation-induced Tie2 degradation upon expression of COMP-Ang1 (Lane 15 and 21). In parallel, distinct Tie2-specific bands at approximately 130 and 95 kDa became detectable (Figure 3B, Panel 2, Lane 5, 11, 17, 23; red arrowheads), indicating/suggesting? destabilization and proteolytic processing of the endogenous Tie2 ectodomain N-terminal of the Tie2 antibody targeting the region around R580. Moreover, additional high molecular weight complexes of phosphorylated myr-Tie2* were detected at 130kDa (Figure 3C, Lane 5, 11, 17, 23, green arrowheads) possibly representing myr-Tie2*-containing, SDS-insoluble, cholesterol-rich lipid rafts.

**Figure 3:**
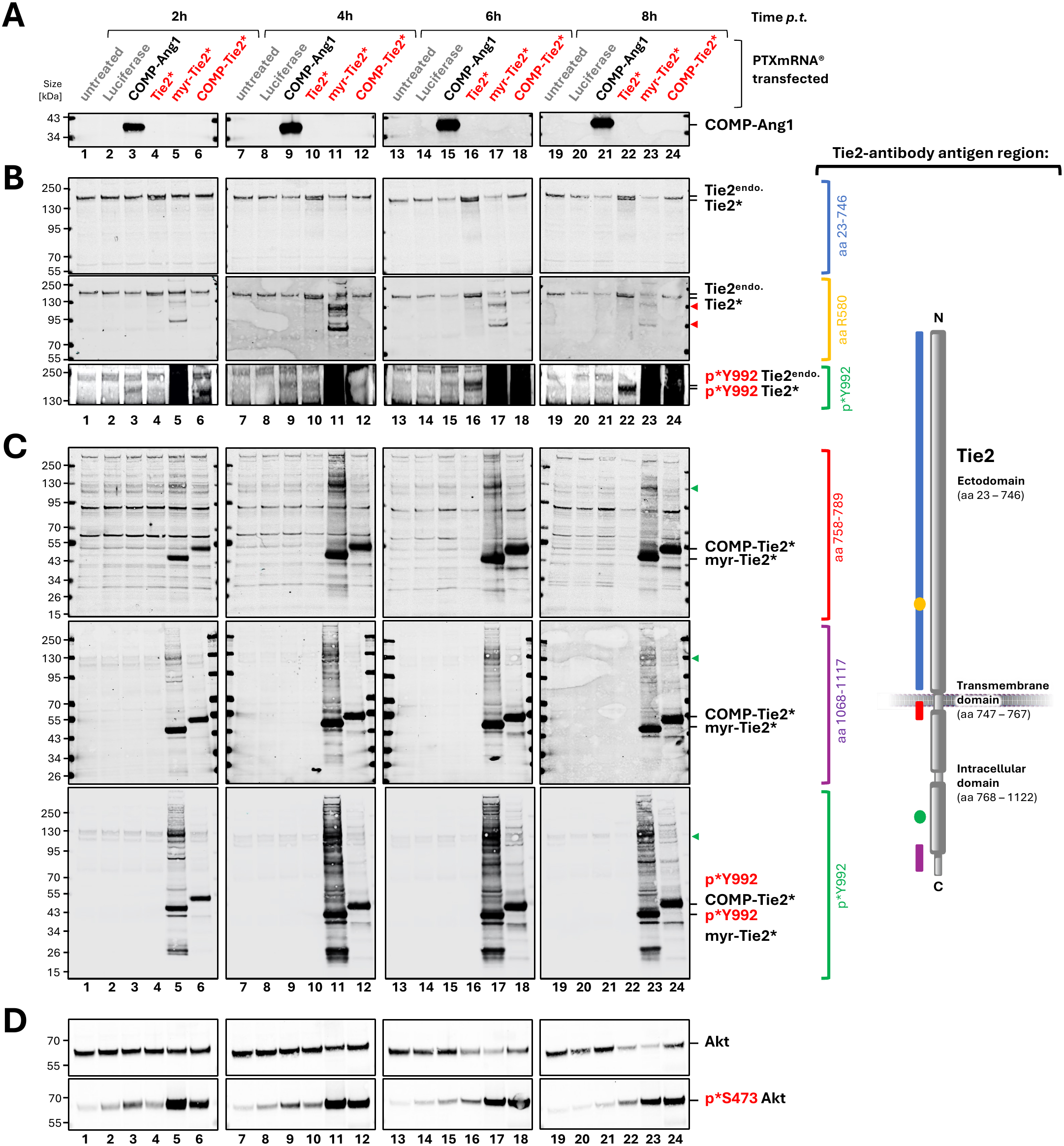
Tie2 pathway activation efficacy of short Tie2* variants in comparison to COMP-Ang1 and full-length Tie2* in HPMECs in vitro. Comparison of time-dependent expression and signaling activity of full-length and engineered Tie2 constructs and COMP-Ang1 in HPMECs. Serum-starved cells were transfected with PTXmRNA® encoding COMP-Ang1, Tie2*, myr-Tie2*, COMP-Tie2* or Luciferase and cells were harvested at indicated time points. Expression of **(A)** COMP-Ang1, and Expression and Tie2 activation (p*Y992) of **(B)** full length Tie2 and **(C)** truncated Tie2 variants were analyzed by SDS-PAGE/Western blotting. **(D)** Downstream Akt expression and phosphorylation (p*S473) levels were assessed as an indicator for Tie2 pathway activation. Red arrow heads indicate endogenous Tie2 ectodomain cleavage fragments; green arrow heads indicate high MW myr-Tie2* complexes. Right, relative location of antigen regions of Tie2 specific antibodies.

These findings demonstrate that both myr-Tie2* and COMP-Tie2* enable faster and more potent Tie2 signaling compared to both ligand-based endogenous Tie2 activation and full-length autoactivating Tie2*. Notably, due to endogenous Tie2 degradation induced by high myr-Tie2* expression, all subsequent studies were conducted using COMP-Tie2*.

### LNP-formulated COMP-Tie2* mRNA enables activation of the canonical Tie2–Akt– FOXO1 signaling axis in vitro

To subsequently examine the Tie2 pathway activation of COMP-Tie2 *in vivo*, COMP-Tie2* mRNA was formulated with LNP002, a cationic LNP designed for RNA delivery to pulmonary vascular endothelial cells^44^. We first confirmed the temporal dynamics of Tie2 pathway activation *in vitro* using COMP-Tie2* in a time-dependent expression study in comparison with a negative control (mRNA-Luciferase) using commercially available transfection reagent LF Messenger Max (Supplementary Figure 2). Following transfection of serum-starved HPMECs, Tie2 phosphorylation at Y992, Y1102, and S1119 became prominent between 4 and 8 hours, decreased by 24 hours, and was undetectable after 48 hours (Figure 4A, Panel 2-4). No decline in endogenous Tie2 levels was observed (Figure 4A, Panel 1). Consistent with Tie2 activation dynamics, downstream Akt and FOXO1 phosphorylation exhibited a comparable temporal kinetic profile, persisting up to 24 hours and returning to baseline by 48 hours (Figure 4B).

**Figure 4.**
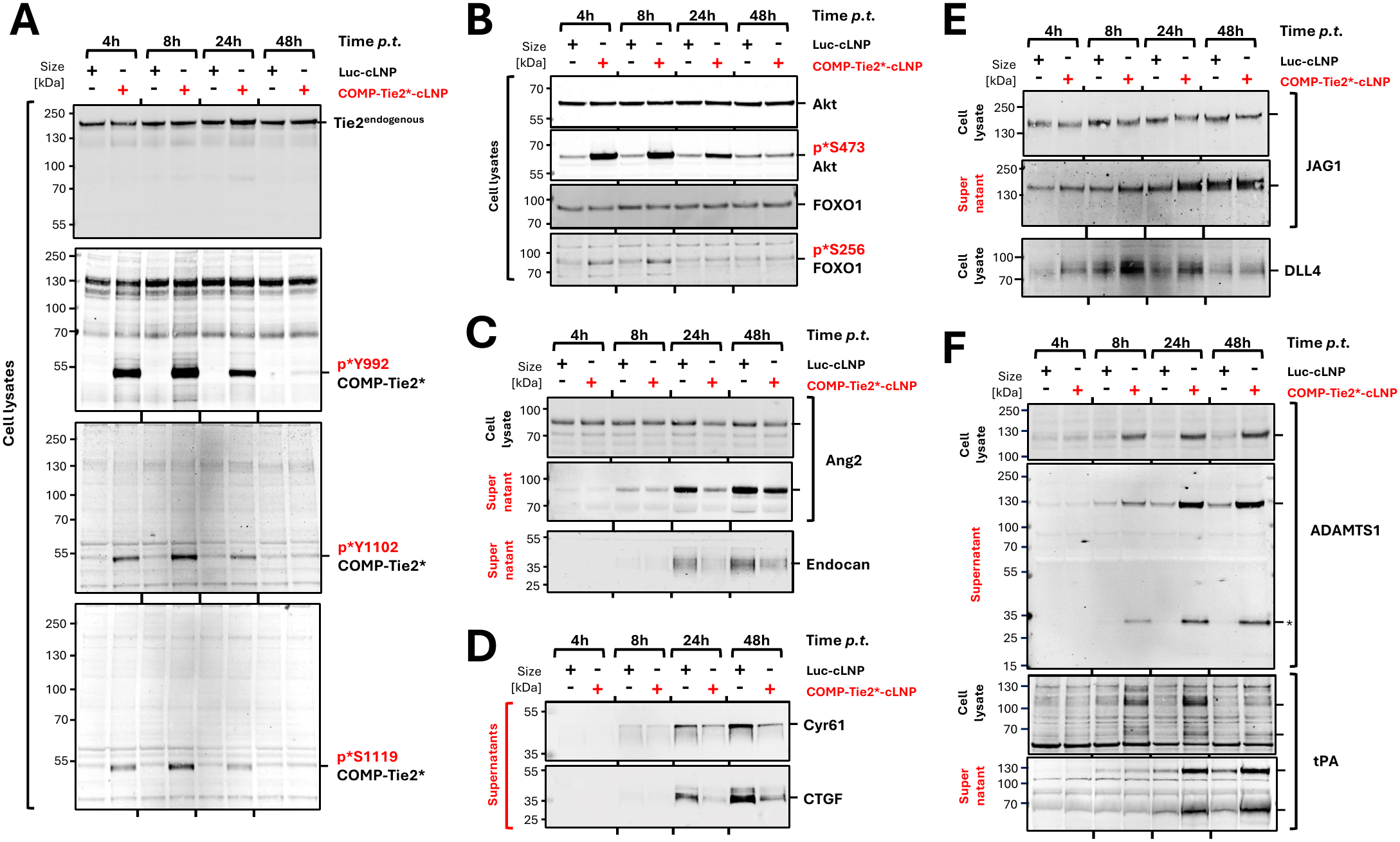
Regulation of endothelial stabilization factors by COMP-Tie2 in vitro. Time-dependent activation of Tie2 signaling and regulation of secreted Tie2 and/or FOXO1-dependent downstream targets following transfection of cationic LNP-formulated COMP-Tie2* mRNA (or Luciferase mRNA as control) in HPMECs. Serum-starved HPMECs were transfected with for 3h in serum-free media and cell lysates and supernatants were analyzed at indicated time points for **(A)** Tie2 phosphorylation and **(B)** Akt–FOXO1 signaling. Protein levels of selected regulators of **(C)** endothelial inflammation (Ang2, Endocan), **(D)** angiogenesis, vascular remodeling and repair (Cyr61, CTGF), **(E)** Notch-based patterning and endothelial cell fate-specification (JAG1, DLL4) and **(F)** vascular microenvironment-modulation (Adamts1, tPA). *, ADAMTS1 Prodomain

To explore Tie2-dependent regulation of secreted proteins, we analyzed selected downstream targets previously reported to be transcriptionally regulated by Tie2-and/or FOXO1-dependent pathways^51,52^. COMP-Tie2* mediated Tie2 signaling includes reduction of pro-inflammatory factors Ang2 and Endocan (Figure 4C). Interestingly, differences in Ang2 levels were more distinct in secreted compared to cellular Ang2 levels (Figure 4C, Panel 1 and 2, Lane 5 vs. 6), suggesting a direct inhibition of Weibel-Palade bodies secretion. In addition, we also observed reduced levels of extracellular matrix-associated, pro-fibrotic factors Cyr61 and CTGF (Figure 4D), along with increased expression of the anti-angiogenic ligand DLL4 and enhanced ectodomain shedding of the pro-angiogenic ligand JAG1 at 24 hours (Figure 4E, Panel 2). In contrast, analysis of ADAMTS1 expression revealed a time-dependent increase of intracellular protein with a peak at 24 hours, followed by reduction at 48 hours, whereas secreted ADAMTS1 and activated ADAMTS1 (indicative of pro-domain cleavage) progressively accumulated in the supernatant over time (Figure 4F, Panel 1 and 2; Supplementary Figure 3A and B). Interestingly, other members of the ADAMTS protein family previously reported to be regulated by the Tie2/Foxo1 pathway at the transcript level, such as ADAMTS4 and ADAMTS5, were no upregulated in response to COMP-Tie2* expression (Supplementary Figure 3D). In addition to ADAMTS1, COMP-Tie2* induced upregulation of anti-thrombotic Tissue-type plasminogen activator (tPA) possessing a very similar kinetic to ADAMTS1, was observed, appearing as a monomer (at around 70 kDa) and higher molecular weight complex (between 100 and 130 kDa) in lysates and supernatant. (Figure 4F, Panel 3 and 4). These data demonstrate coordinated Tie2 signaling-dependent regulation of secreted and membrane-anchored endothelial mediators associated with vascular stabilization. This indicates that COMP-Tie2* induces sustained yet transient activation of the canonical Tie2–Akt–FOXO1 signaling cascade and promotes functional expression of FOXO1-dependent downstream targets.

In addition to changes in gene expression, Tie2 activity has been shown to modulate adherens junctions by regulation of VE-Cadherin stability. Consequently, immunofluorescence microscopy of VE-Cadherin in COMP-Tie2*-transfected HPMECs showed increased VE-Cadherin-positive junctional membrane overlaps consistent with the role of Tie2 in VE-Cadherin stabilization and permeability regulation^53,54^ (Supplementary Figure 3).

### *In vivo* activation of Tie2 signaling following systemic mRNA delivery with cLNP targeting the lung vasculature

Having established robust Tie2 pathway activation *in vitro* after transfection of LNP-formulated COMP-Tie2* mRNA *in vitro*, we next evaluated COMP-Tie2* activity *in vivo*. We have previously described the application of cationic LNPs targeting the lung vascular endothelium in mice for positioning and expression of therapeutic COMP-Ang-1 mRNAs^44^ (Dunne et al., in Preparation). The same cLNP formulation was employed here to deliver the COMP-Tie2* mRNA into the lung tissue after systemic administration in mice. Mice were intravenously injected (bolus tail vein) with COMP-Tie2*-cLNP (0.75 mg/kg), and tissues were collected at defined time points (Figure 5A). Phosphorylated COMP-Tie2* (p*Y992) was clearly detected in lung lysates at 2 hours, with peak signal at 4 hours post-administration (Figure 5A; Supplementary Figure 4). In contrast, no p*Y992 signal with a COMP-Tie2*-corresponding size was detectable in liver, spleen, heart, or kidney samples, demonstrating lung-selective expression following systemic delivery (Figure 5B; Supplementary Figure 4).

**Figure 5.**
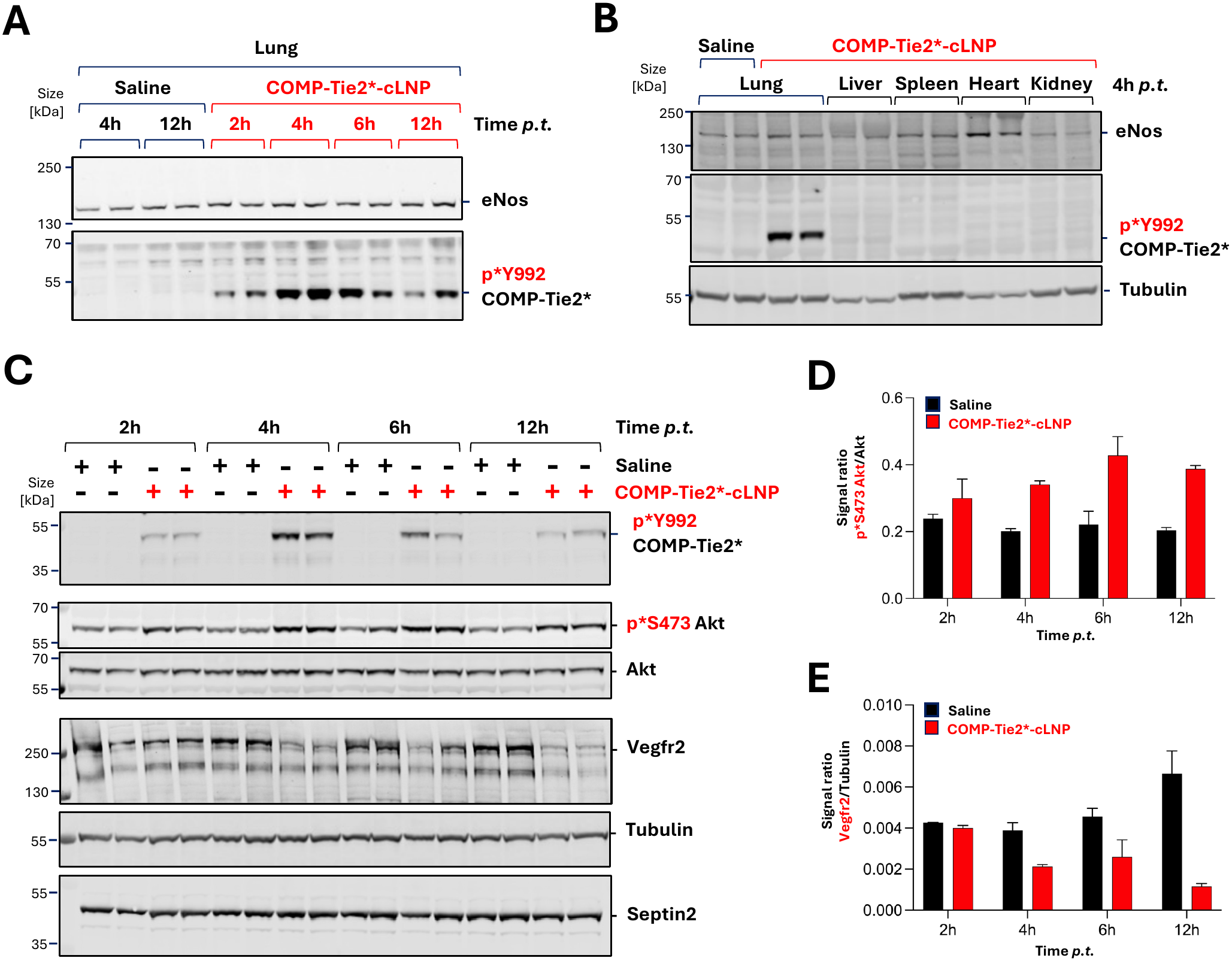
Lung-selective expression and activation of COMP-Tie2 following systemic administration in vivo. **(A)** Time-dependent expression of COMP-Tie2* in murine lung tissue of individual animals following intravenous administration of LNP-formulated COMP-Tie2* mRNA in mice (0.75 mg/kg). **(B)** Tissue distribution analysis of COMP-Tie2* expression 4 hours after LNP administration using phospho-Tie2 (Y992)–specific antibodies. Endothelial nitric oxide synthase (eNOS) served as endothelial loading control. **(C)** Western blot analysis of whole lung tissue lysates from individual animals demonstrating time-dependent COMP-Tie2* phosphorylation, Akt activation and VEGFR2 reduction following intravenous administration of COMP-Tie2*-cLNP. Akt served as loading control for Akt phoisphorylation and Septin served as loading control for endothelial cell content. **(D)** Quantification of Akt activation and **(E)** VEGFR2 reduction in whole lung tissue expressed as pS473 Akt/Akt signal ratio. Data are shown as mean ± SEM.

To assess downstream signaling, lung lysates were further analyzed for Akt activation. Although endothelial cells account for only about 30-50% of total lung tissue^55–57^, a clear increase in Akt phosphorylation was observed compared with saline-treated controls and persisted for up to 12 hours post-injection (Figure 5C, D). In parallel, COMP-Tie2* expression led to a time-dependent reduction of VEGFR2 protein levels in lung tissue (Figure 5C, E).

Together, these data demonstrate that systemic delivery of COMP-Tie2* via mRNA/cLNP enables selective activation of Tie2 signaling in the pulmonary endothelium *in vivo* and represents a promising pharmacological means to promote endothelial stabilization towards improving microvacular barrier function.

## Discussion

A major characteristic in inflammatory vascular diseases such as ARDS is the profound dysregulation of endogenous Tie2 signaling. Angiopoietin-1 (Ang1) and Angiopoietin-2 (Ang2) are counteracting ligands for the Tie2 receptor, primarily regulating vessel stability during inflammation^58^. An imbalance in the Ang2/Ang1 ratio is a critical marker for disease severity in conditions like sepsis, acute lung injury, and diabetic retinopathy. Ang2 binds to the same site on the Tie2 receptor as Ang1 but typically fails to activate it as effectively, thereby antagonizing Ang-1-mediated endothelial stabilization^15,59^. Elevated circulating Ang2 levels strongly correlate with disease severity and mortality^60–62^. Compounding this, inflammatory protease activity promotes ectodomain shedding of Tie2^36,38,63,64^, reducing functional receptor availability and generating soluble Tie2 (sTie2) fragments that further sequester Ang1. Under these inflammatory conditions, strategies aimed at increasing agonistic Tie2 ligand levels—either through recombinant protein administration or mRNA-mediated Ang1 expression—are intrinsically limited by reduced receptor accessibility.

The COMP-Tie2* variant described here directly overcomes these limitations by bypassing the extracellular ligand–receptor interface. Because receptor activation is encoded within the engineered Tie2 molecule itself (Figure 1A-C), signaling is insensitive to Ang2 competition and does not depend on intact endogenous Tie2 ectodomains (Figure 1D, E). Taken together our data demonstrates that ligand-independent receptor engineering represents a mechanistically distinct strategy for therapeutic Tie2 activation. By eliminating the need for extracellular ligand binding, membrane-anchored short Tie2 variants circumvent multiple counteracting mechanisms, including endogenous regulatory and inhibitory processes such as inflammation-induced receptor shedding. These endogenous constraints have limited the efficacy of previous strategies, including recombinant Ang1-based therapies^21–23,25,28^ , agonistic Tie2 antibodies^26, 65^or Ang2-neutralizing approaches, such as Ang-2 capturing antibodies^66,67^ .

Productive Tie2 signaling depends not only on kinase activity but also on precise spatial organization of the receptor within the endothelial plasma membrane^68–70^. Physiological Tie2 activation by Ang1 relies on ligand-mediated receptor clustering at endothelial cell–cell junctions, where Tie2 interfaces with adherends junction complexes and cytoskeletal regulators that govern vascular barrier integrity^71–73^.

To enforce plasma membrane localization by posttranslational myristylation within cholesterol-enriched junction-associated membrane microdomains^74–76^, a fusion of an N-terminal Src myristylation domain with a constitutively active intracellular Tie2 domain (myr-Tie2*) was designed (Figure 2A). The same principle of membrane targeting has been successfully demonstrated to enforce FGFR signaling induction^77,78^ by using mutated and myristylated FGFR intracellular kinase domains. In a parallel experimental approach, incorporation of the COMP pentamerization domain to the transmembrane and constitutively active intracellular Tie2 domain (COMP-Tie2*) enabled higher-order receptor clustering^21,79^, functionally mimicking Ang1-induced Tie2 oligomerization while avoiding ligand dependency.

In line with this design, myr-Tie2* and COMP-Tie2* induced faster and more robust Tie2 autophosphorylation than COMP-Ang1 or full length Tie2*, and resulted in stronger downstream signaling (Figure 3D). Surprisingly, myr-Tie2* expression resulted in endogenous Tie2 ectodomain degradation (Figure 3B, red arrowheads) and morphological changes towards spindle-like appearance and cell detachment (not shown), despite high downstream signaling activity, highlighting the necessity of precise spatial control of Tie2 membrane localization^73^.

Beyond direct junctional effects (Supplementary Figure 3), COMP-Tie2* induced broader endothelial reprogramming. Sustained activation of the PI3K-Akt pathway^80^ resulted in phosphorylation and inhibition of the transcription factor FOXO1^81,82^, a central regulator of endothelial activation and vascular permeability (Figure 2D, 4B, Supplementary Figure 2B). Consistent with this, COMP-Tie2* promoted expression of FOXO1-regulated vascular-stabilizing mediators including ADAMTS1^51,83^ (Figure 4F, Supplementary Figure 2B) and tissue plasminogen activator^51,84,85^ (tPA, PLAT) (Figure 4B), while repressing multiple endothelial destabilization factors such as Ang2, Endocan (ESM1), connective tissue growth factor (CTGF, CCN2), and cysteine-rich angiogenic inducer 61 (Cyr61, CCN1) (Figure 4C, D). Although several of these genes have previously been identified as Tie2-regulated, primarily at the transcript level via FOXO1 inhibition^86^ or modulation of additional transcription factors such as KLF2^87^, the present study demonstrates their coordinated regulation at the protein level, underscoring the functional relevance of sustained pathway activation. Consistent with previous reports^88^, we observed a significant increase in endothelial DLL4 expression upon COMP-Tie2 expression (Figure 4E). Remarkably, this DLL4 upregulation was accompanied by enhanced JAG1 cleavage. In the established endothelial “tip-stalk” hierarchy, DLL4 is primarily secreted by leader tip cells as to activate Notch signaling in adjacent cells, thereby suppressing the sprouting phenotype and inducing a stable stalk cell identity^89^ with increased cell-cell adhesion. Conversely, JAG1 typically functions as a pro-angiogenic antagonist to DLL4. JAG1 attenuates Notch signaling in stalk cells to facilitate further branching and tip cell formation by competing for Notch receptors ^90,91^. Collectively, the concurrent upregulation of DLL4 and the increased proteolytic shedding of JAG1, driven by COMP-Tie2*, suggest a decisive shift toward vascular stabilization. By maximizing DLL4-mediated Notch activation while simultaneously removing the antagonistic influence of JAG1, this mechanism robustly reinforces the stalk cell phenotype and prevents excessive, unproductive vessel sprouting^92^. Consistent with this observation, we detected enlarged VE-cadherin-positive regions at cell-cell-junctions, indicating enhanced intercellular adhesion as previously reported^53^ (Supplementary Figure 3).

ADAMTS1 has been implicated in extracellular matrix remodeling and in limiting pathological vascular permeability^93^. It exerts anti-angiogenic effects by directly binding and sequestering VEGF165 (vascular endothelial growth factor), thereby preventing its interaction with its receptor (VEGFR2, KDR), and significantly inhibiting VEGFR2 phosphorylation, which in turn suppresses endothelial cell proliferation^94,95^. Induction of Adamts1 expression by the Ang1-Tie2-Foxo1 cascade has been shown previously^83,96,97^. Like Adamts1, tPA has been shown to be a Tie2-responsive gene^84,98^. Increased endothelial tPA expression is associated with preservation of microvascular patency and limitation of fibrin deposition, processes frequently disrupted in acute lung injury and ARDS^99,100^. Conversely, Ang2 is a well-established antagonist of Tie2 signaling and a clinical biomarker of disease severity in ARDS^101,102^. Suppression of Ang2 therefore represents a key functional consequence of sustained Tie2 activation. Similarly, the downregulation of endocan, a pro-inflammatory, pro-angiogenic factor and established marker of endothelial dysfunction^103,104^, along with CTGF and Cyr61, key regulators of angiogenesis and vascular remodeling during wound healing and inflammation, suggests attenuation of endothelial activation, vascular remodeling, and inflammatory amplification^105–108^.

A distinct advantage of this novel mRNA-cLNP platform is its ability to act in a highly spatially restricted manner, targeting the endothelium specifically at the affected vascular bed of lung injury. Importantly, LNP-based mRNA delivery also enables transient receptor expression as previously shown^44^. Following systemic administration, COMP-Tie2* expression delivered via lung-endothelium targeted lipid nanoparticles LNP002, was selectively restricted to pulmonary tissue, leading to localized, ligand-independent activation of the Tie2 pathway without detectable activation in peripheral organs (Figure 5B; Supplementary Figure 4). This precision enables rapid activation of the Tie2 pathway and provides superior pharmacodynamic properties compared to recombinant protein therapies, which often require continuous infusion because of their rapid systemic clearance. In addition, COMP-Tie2* expression in murine lung endothelium resulted in a time-dependent activation of Akt and a reduction of VEGFR2 protein levels (Figure 5C, D). Regulation of VEGFR2 expression has been shown through FOXO1 inhibition and through DLL4 activation^88,109–115^, two processes occurring upon COMP-Tie2* expression. In addition, Tie2 activation has been demonstrated to recruit IQGAP1, which is typically bound to VEGFR2, towards Rac1, thereby stabilizing VE-Cadherin and downregulating IQGAP1, thus reducing VEGFR2 stability^116–119^. This process likely reflects a shift toward an endothelial quiescent phenotype characterized by reduced VEGF responsiveness, which may be advantageous in inflammatory lung disease where excessive VEGFR2 signaling may contributes to vascular leakage^120^.

Although the COMP-Tie2 construct enables robust, ligand-independent signal activation, its expression duration remains comparatively short. This is due to continuous receptor phosphorylation, which naturally recruits ubiquitin ligases such as Cbl. Cbl subsequently ubiquitinates COMP-Tie2, initiating its rapid internalization and cellular degradation^43^. To counteract this limitation and to further extend the receptor’s half-life, future modifications could introduce targeted mutations at the specific lysine residues required for ubiquitination.

Preventing this ubiquitination step would delay receptor breakdown and significantly prolong COMP-Tie2* signaling activity.

Building upon our previous research, an intriguing future perspective is the potential combinatorial delivery of the ligand-independent COMP-Tie2 construct alongside ligand-based Tie2 agonists. We recently demonstrated that the intravenous administration of cationic LNPs formulated with mRNA encoding the hyperactive fusion protein COMP-Ang1 selectively targets capillary endothelial cells of the lung. This localized, mRNA-directed expression of COMP-Ang1 effectively attenuated inflammation-induced vascular leakage and reduced neutrophil infiltration in murine models. Given that both the COMP-Tie2* and COMP-Ang1 approaches utilize the identical, highly specific cLNP delivery platform, multiplexing these mRNAs into a single therapeutic formulation represents a highly feasible strategy. Such a combined approach could yield profound synergistic effects in severe inflammatory conditions such as ARDS. While COMP-Tie2* ensures a robust, sustained baseline of endothelial stabilization that is fully resistant to Ang2 antagonism and inflammation-induced receptor shedding, locally secreted COMP-Ang1 can simultaneously act in both paracrine and autocrine modes to activate remaining intact endogenous Tie2 receptors on neighboring cells.

In conclusion, our findings establish COMP-Tie2* as a next-generation modulator of the Tie2 pathway that overcomes key limitations of ligand-based strategies. By decoupling receptor activation from extracellular angiopoietin regulation, COMP-Tie2* enables sustained endothelial stabilization even under conditions of high Ang2 burden and potential Tie2 shedding.

Looking ahead, a dual-action approach combining COMP-Ang-1 with COMP-Tie2* could harness the complementary strengths of ligand-mediated and receptor-intrinsic activation, thereby maximizing comprehensive barrier protection supporting its potential use in inflammatory vascular diseases such as ARDS. Moreover, COMP-Tie2* may represent a novel, mechanistically distinct anti-angiogenic strategy in oncology: while current agents, such as Ang2/VEGF-A bispecific antibodies, act by neutralizing extracellular angiogenic drivers, COMP-Tie2* directly enforces intracellular Tie2 activation, counteracting Ang2 expression/secretion and VEGF signaling at their source.

## Materials & Methods

### mRNA synthesis

Codon-optimized, CleanCap™-equipped mRNAs containing 5’ UTR sequence and signal peptide sequence of human MCP1 (NM_002982.3), 3’ UTR sequence of human vWF (NM_000552.4) and coding regions of COMP-Ang1, Tie2, Tie2 R849W, Tie2 L914F, Tie2 849W/L914F (Tie2*), myr-Tie2*, COMP-Tie2* and Firefly Luciferase and 120nt PolyA tail were synthesized and purified by BioSpring GmbH (Frankfurt am Main, Germany) and AmpTec (now Merck, Hamburg, Germany). Uridine was globally replaced with N1-methylpseudouridine.

### LNP preparation and characterization

For manufacturing LNP002 the cationic lipid (L-Arginyl)-L-2,3-diamino propionic acid-N-palmityl-N-oleyl-amide, Pantherna Therapeutics, Hennigsdorf, Germany), the helper lipid 1,2-Diphytanoyl-sn-glycero-3-phosphoethanolamine (Corden Pharma, Plankstadt, Germany) and the PEGylated lipid 1,2-distearoyl-sn-glycero-3-phosphoethanolamine-N-[methoxy(polyethylene glycol)-2000] (ammonium salt) (Avanti Polar Lipids, Alabaster, USA) were dissolved in pure ethanol with a molar ratio of 50:49:1, mixed with mRNA dissolved in 10 mM sodium citrate buffer at pH 5.5 resulting in a lipid:mRNA mass ratio of m:m=20, with mRNA concentration of 100–280 μg/ml in the final formulation. Specifically, the aqueous mRNA solutions and the ethanolic lipid mixtures were combined at a volume ratio of 2:1 (aqueous:ethanol) and flow rates of 18 ml/min using a microfluidic mixer (NanoAssemblr®; Precision Nanosystems, Vancouver, Canada). Formulations were dialyzed against 10 mM Tris (pH 7.4) with 9 % sucrose in Slide-A-Lyzer dialysis cassettes (MWCO 3.5k, Thermo Scientific, Rockford, IL, USA) for at least 18 h at 4 °C. The formulations were tested for particle size and zeta potential (Zetasizer Nano ZS, Malvern Instruments, Malvern, UK) as well as for RNA encapsulation and for total RNA content (Quant-iT RiboGreen RNA Assay Kit, Thermo Fisher Scientific, Waltham, USA). The formulations were found to have a Z-Average between 80 and 90 nm, a PDI < 0.1 and a Zeta potential between +5-10 mV using 10 mM Tris (pH 7.4) with 9 % sucrose as suspension buffer. The mRNA encapsulation was always > 95 %. The formulations were stored at –80 °C until further use.

### Cell transfection in vitro

Hela cells were transfected with mRNAs using Lipofectamine MessengerMax (Invitrogen, cat. #LMRNA001) according to manufacturer’s protocol. Briefly, cells were seeded in 150 cm^2^ culture flasks (TPP, cat. #90151) at 50 % confluency in serum-free DMEM (Gibco, cat. #61965-026). 15 h later, cells were transfected for 2 h with indicated amounts of LF MessengerMax-formulated mRNAs according to manufacturer’s protocol. Transfection media was removed and replaced with fresh serum-reduced DMEM media for indicated time points. Human pulmonary microvascular endothelial cells (HPMEC) (Promocell, cat. #C-12281) were seeded onto 6-well plates (Thermo Scientific, cat. #140675) and left for 3 days until 100 % confluency in full supplemented ECM MV2 cell media (Promocell, cat. #C-22022). 15 hours before transfection cells were placed in serum-reduced ECM MV2 media (containing 10 % of original cell growth supplement MV2). HPMEC were transfected with preformulated mRNA-cLNP at indicated concentrations in serum-free ECM media for 2 h. After transfection, media was replaced with serum-reduced ECM MV2.

### Western Blot analysis

Cells were washed with ice-cold phosphate buffered saline (PBS) twice and lysed with ice-cold RIPA buffer (Thermo Scientific, cat. #89901) supplemented with Halt™ Protease and Phosphatase Inhibitor Cocktail (Thermo Scientific, cat. #78441). The supernatants were removed and tenfold concentrated using Amicon 3 kDa spin columns (Merck, cat. #UFC900308) and then mixed with 4xSDS-loading buffer for subsequent Western blot analysis.

Tissue samples were solubilized in T-PER™ Tissue Protein Extraction Reagent (Thermo Scientific, cat. #78510) supplemented with Pierce Protease and Phosphatase Inhibitor tablets (Thermo Scientific, cat. #A32958) using Qiagen TissueLyser LT (Qiagen, cat. #69980). A mixture of supernatant, cell or tissue lysate, 50 mM dithiothreitol NuPAGE reducing agent (Invitrogen, cat. #NP0009) and NuPAGE sample buffer (Invitrogen, cat. #NP0007) was heated at 70 °C for 10 min, electrophoresed in NuPAGE 4–12 % Novex Bis-Tris Gels and MOPS buffer (all from Invitrogen, cat. #NP0335BOX, NP0001) and transferred to nitrocellulose membrane (Cytiva, cat. #10600003; NuPAGE™ Transfer Buffer (Invitrogen, cat. #NP00061). After a 60 min blocking step (Intercept® Blocking Buffer; LiCor, cat. #927-60001) membranes were washed with Tris Buffered Saline with Tween® 20 (CST, cat. #9997S) and probed at 4 °C overnight with specific primary antibodies diluted in Intercept Antibody Diluent (LiCor, cat. #927-65001). Binding of primary antibodies was detected by near infrared IRDye secondary antibodies (LiCor, cat. #926-32213, 926-32214, 926-68072) and near infrared signals were analyzed using LiCor Odysee CLx. Band intensities were analyzed using Empiria Studio Software 2.1.0134 (LiCor). Primary antibodies used were: Angiopoeitin-1 (abcam, cat. #ab183701), Angiopoeitin-2 (abcam, cat. #ab8452), N-terminus specific Tie2 (CST, cat. #7403), N-terminus specific Tie2 (R&D Systems, cat. # MAB313), C-terminus specific Tie2 (Merck, cat. #SAB4502942), C-terminus specific Tie2 (NSJ Bioreagents, TEK/9412 cat.#V6010), pY992 Tie2 (R&D Systems, cat. #AF2720), pY1102 Tie2 (R&D Systems, cat. #AF3909), pS1119 Tie2 (CST, cat. #4226), Akt (CST, cat. #9272 and #2920), pS473 Akt (CST, cat. #4060), S6K (CST, cat. #9202), pT389 S6K (CST, cat. #9234), FoxO1 (CST, cat. #2880), pT24 FoxO1 (CST, cat. ##9464), pT389 FoxO1 (CST, cat. #9461), Adamts1 (CST, cat. #12897), CTGF (CST, cat. #86641), Cyr61 (CST, cat. #14479), DLL4 (CST, cat. # #96406), Jag1 (R&D Systems, cat. #AF1277), VEGFR2 (CST, cat. #2479), Endocan/ESM1 (R&D Systems, cat. #AF1810), tPA (R&D Systems, cat. #AF7449), eNOS (CST, cat. #32027), Septin2 (Merck, cat. #HPA018481).

### Fluorescent microscopy

HPMEC were seeded in Chamber Slides (NUNCTM Lab-TekTM, cat. #154534PK) at 1x10^5^ cells/ml in endothelial MV2 growth medium. Cells were incubated for 3-5 days at 37 °C in 5 % CO_2_. Cells were washed 2 times with pre-warmed PBS (Thermo Scientific, cat. #14190), final PBS wash was replaced with endothelial MV2 basal medium devoid of serum and supplements. Formulations containing mRNA were diluted in basal medium and added to each well for a final concentration of 0,38 ng/well. Cells were returned to the incubator for an additional 2 hours of incubation. Media containing formulations was removed and replaced with 10 % MV2 growth medium for 16 h. Cells were fixed in 4 % PFA (Thermo Scientific, cat. #15670799) for 10 min at room temperature (RT) and subsequently permeabilized in PBS containing 0.02 % Triton-X (PBS-T) for 2x 10 min washes on a rocker at RT. Primary antibody mix was made up with rabbit anti-VE-Cadherin (CST, cat. #2500) diluted at 1:400 in 2 % donkey immunobuffer containing 0.02 % Triton-X, rocking at 4 °C overnight. Primary antibodies were removed and washed in PBS-T 2x 10 min on a rocker at RT. Secondary antibody mix was made up with donkey anti-rabbit (H+L) highly cross-adsorbed Alexa FluorTM 647 (Thermo Scientific, cat. #A-31573) diluted at 1:400 and Alexa FluorTM 594 Phalloidin (Thermo Scientific, cat. #A12381) at 1:2000 in 2 % donkey immunobuffer containing 0.02 % Triton-X rocking at RT for 5 h. Cells were washed in PBS-T for a further 2x 10 min at RT. Nuclear staining was carried out with Hoechst (Tocris, cat. #5117) diluted at 1:500 in PBS for 15 min, followed by a final PBS-T wash. Fluorescent images were acquired with a Nikon Eclipse Ti-U with an ELWD S Pan Fluor 40X/0.6 objective lens. Images were processed using ImageJ.

### Animals

Female C57BL/6N mice (8 to 10 weeks, 18 to 20 g; Charles River, Sulzfeld, Germany) were used for all experiments. All animal experiments were approved by institutional and governmental German authorities (“Tierschutzbeauftragte” (animal welfare officer) and “Tierschutzausschuss” of Charité–Universitätsmedizin Berlin; and by the Landesamt für Gesundheit und Soziales Berlin; approval ID A0179/20) and were in accordance with the Federation of European Laboratory Animal Science Associations (FELASA) guidelines and recommendations for the care and use of laboratory animals (equivalent to American ARRIVE). Mice were randomly assigned to the appropriate experimental groups and were kept in closed, individually ventilated cages with filter hoods (type II-L, ZOONLAB), under specific pathogen-free conditions, with free access to food and water, room temperature between 20 and 22 °C, air humidity between 50 and 65 % and 12 h light/dark cycle.

## Author contributions

K.R., C.M.D., O.K., and D.T. conducted the experiments and were involved in methodology development and the formal analysis of data. J.K., A.S., and DT conceptualized the experiments and were engaged in project administration.

## Acknowledgements

We thank Diana Behrens from EPO Berlin-Buch GmbH for carrying out the *in vivo* study and Katharina Ahrens for reviewing the maniscript.

---

**Supplementary Figure 1:**
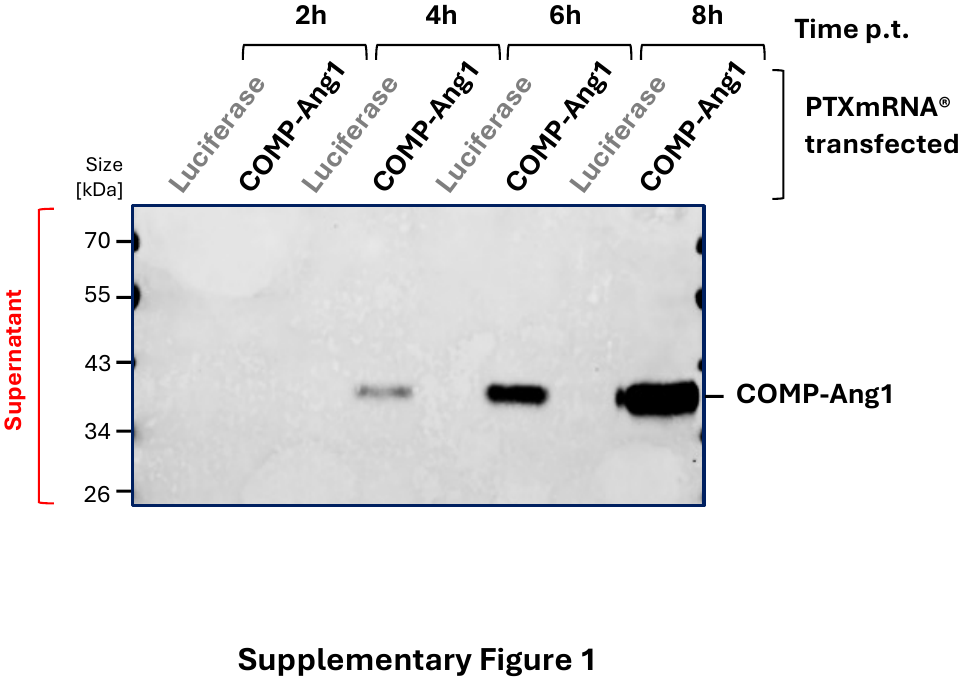
Analysis of COMP-Ang1 secretion. Analysis of COMP-Ang1 secretion at indicated time points into HPMEC supernatant after COMP-Ang1 mRNA transfection in HPMEC supernatant by SDS-PAGE/Western Blotting.

**Supplementary Figure 2:**
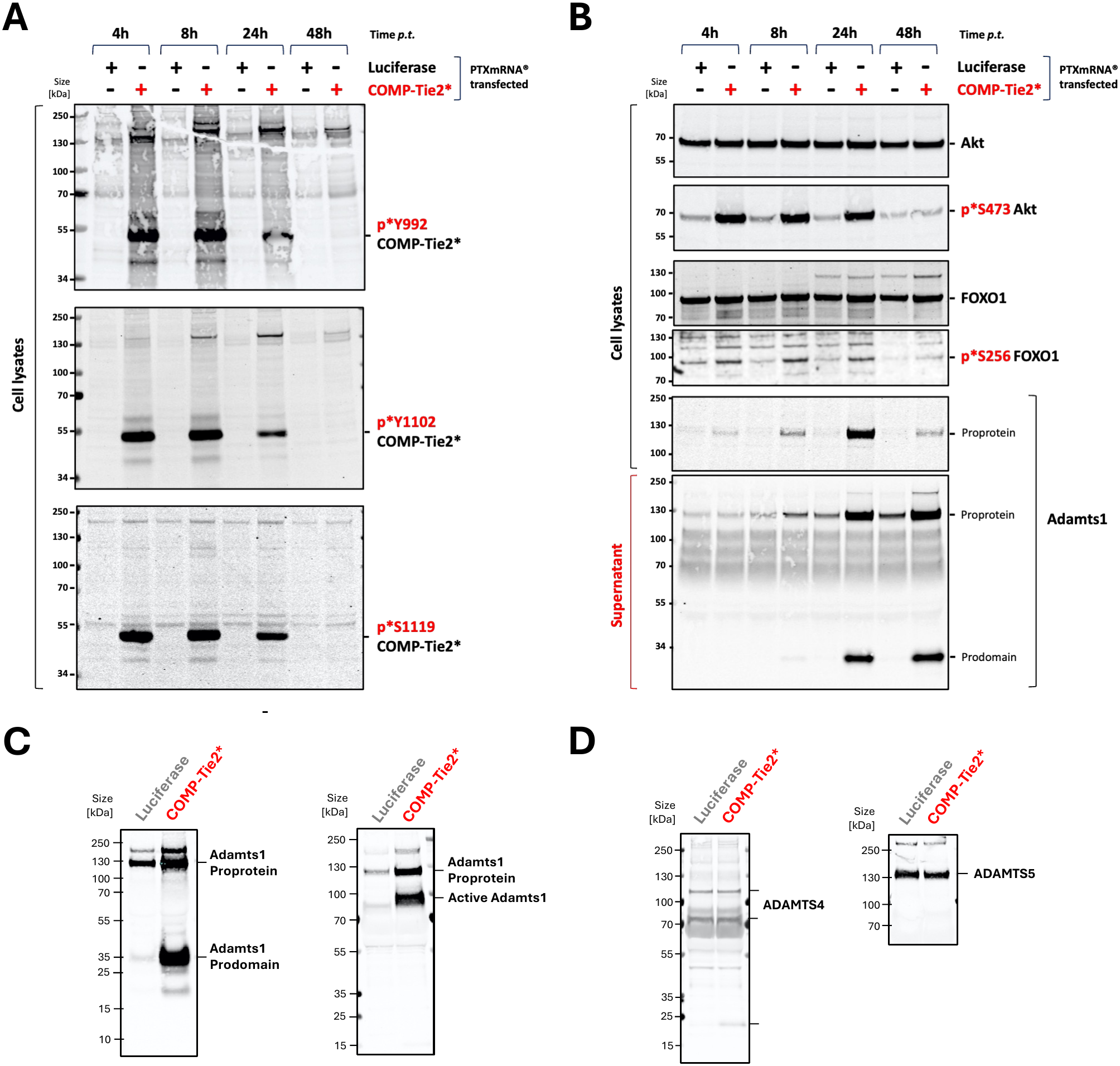
Time-dependent activation of the canonical Tie2–Akt–FOXO1 signaling axis and induction of Adamts1 expression, secretion and activation after MessengerMax-mediated transfection of COMP-Tie2* PTXmRNA in vitro. Time-dependent expression and phosphorylation of COMP-Tie2* in HPMECs. Cells were transfected with LF Messenger Max for indicated time points with of PTXmRNA® encoding COMP-Tie2* or Luciferase using LF Messenger Max. SDS-PAGE/Western blot analysis was performed from protein lysates at indicated time points after transfection and assessed for (A) Tie2 autophosphorylation at Y992, Y1102, and S1119. (B) Tie2 pathway downstream signal transduction was analyzed by Akt activation at S473, FOXO1 inhibition at S256, and induction and secretion of ADAMTS1. Endogenous Tie2, Akt, and FOXO1 served as loading controls. (C) Analysis of Adamts1 secretion and activation in HPMEC cell supernatant 24h after transfection of COMP-Tie2* mRNA by SDS-PAGE/Western blotting using Adamts1 antibodies specific for N-terminal Prodomain (left) and specific raised against prodomain-less, active Adamts1 (right). (D) Analysis of Adamts4 and Adamts5 in HPMEC cell supernatant 24h after transfection of COMP-Tie2.

**Supplementary Figure 3.**
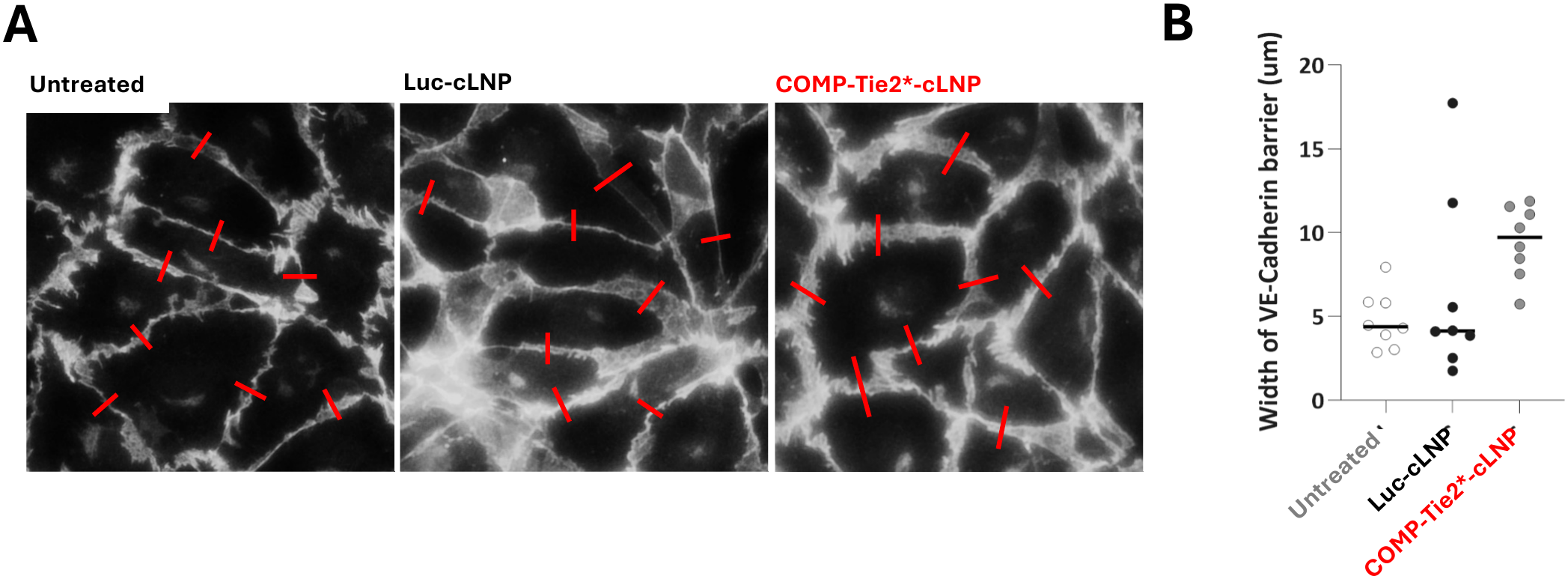
Regulation of endothelial junction integrity by LNP-formulated COMP-Tie2 in vitro. Immunofluorescence analysis of endothelial cell–cell junctions following LNP-mediated COMP-Tie2* expression. HPMECs were stained for VE-cadherin, and junctional width was quantified. Representative images **(A)** and quantitative analysis **(B)** demonstrate increased endothelial barrier thickness following COMP-Tie2* expression compared with untreated or Luciferase-LNP controls.

**Supplementary Figure 4.**
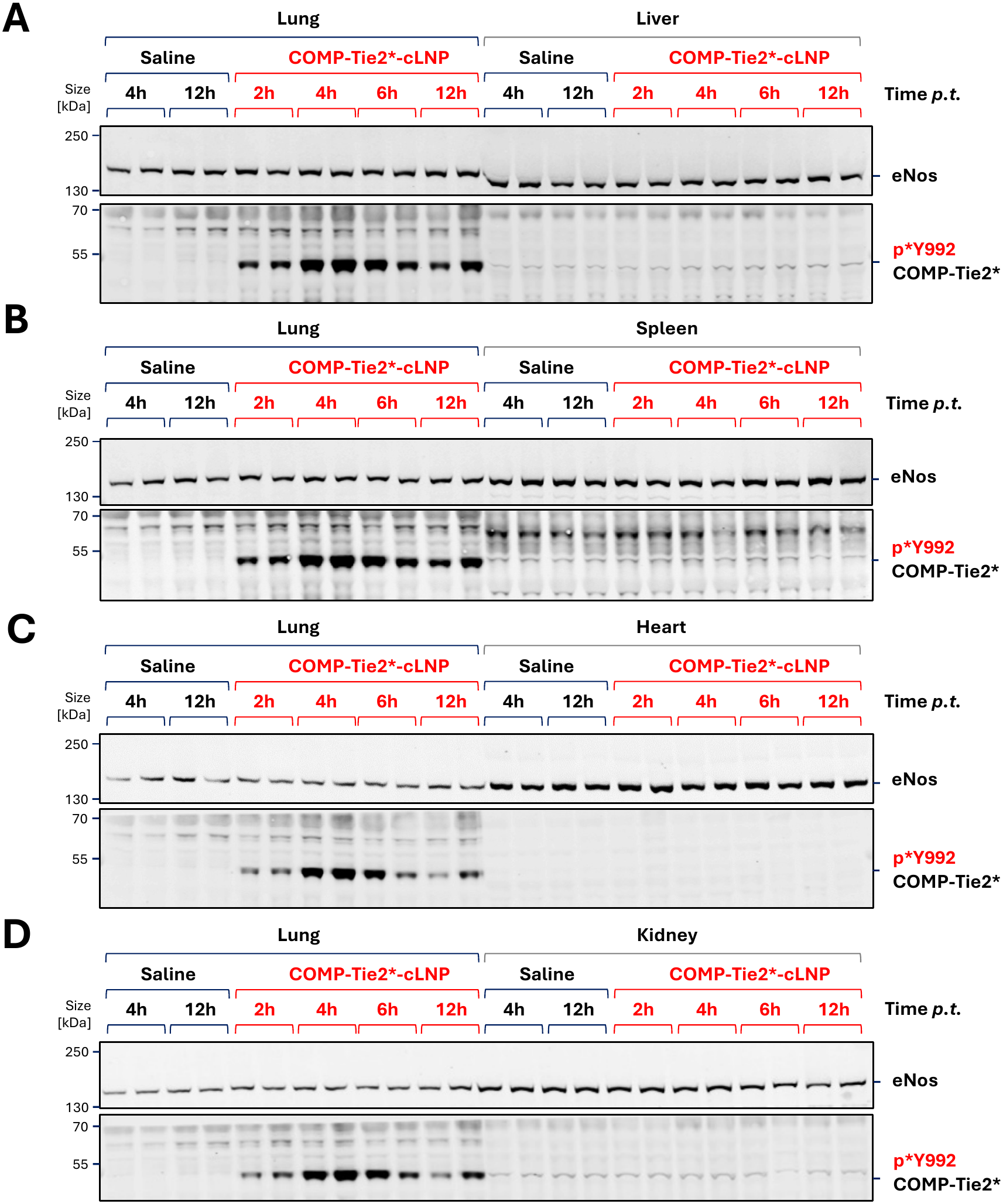
Lung-selective expression and activation of COMP-Tie2 following systemic administration in vivo. **(A–D)** Tissue distribution analysis of COMP-Tie2* expression in individual animals following intravenous administration of LNP-formulated COMP-Tie2* mRNA in mice (0.75 mg/kg). Lung tissue lysates were analyzed in parallel with liver (A), spleen (B), heart (C), and kidney (D) tissues at indicated time points for COMP-Tie2* expression using phospho-Tie2 (Y992)–specific antibodies. Endothelial nitric oxide synthase (eNOS) served as endothelial loading control.

## References

1. Joffre, J., Hellman, J., Ince, C., and Ait-Oufella, H. (2020). Endothelial Responses in Sepsis. Am. J. Respir. Crit. Care Med. 202, 361–370. 10.1164/RCCM.201910-1911TR.

2. Matthay, M.A., Zemans, R.L., Zimmerman, G.A., Arabi, Y.M., Beitler, J.R., Mercat, A., Herridge, M., Randolph, A.G., and Calfee, C.S. (2019). Acute respiratory distress syndrome. Nat. Rev. Dis. Primers 5. 10.1038/S41572-019-0069-0.

3. Van Der Heijden, M., Van Nieuw Amerongen, G.P., Chedamni, S., Van Hinsbergh, V.W.M., and Groeneveld, A.B.J. (2009). The angiopoietin-Tie2 system as a therapeutic target in sepsis and acute lung injury. Expert Opin. Ther. Targets 13, 39–53. 10.1517/14728220802626256.

4. Thurston, G., Suri, C., Smith, K., McClain, J., Sato, T.N., Yancopoulos, G.D., and McDonald, D.M. (1999). Leakage-resistant blood vessels in mice transgenically overexpressing angiopoietin-1. Science 286, 2511–2514. 10.1126/SCIENCE.286.5449.2511.

5. Thurston, G., Rudge, J.S., Ioffe, E., Zhou, H., Ross, L., Croll, S.D., Glazer, N., Holash, J., McDonald, D.M., and Yancopoulos, G.D. (2000). Angiopoietin-1 protects the adult vasculature against plasma leakage. Nat. Med. 6, 460–463. 10.1038/74725.

6. Gamble, J.R., Drew, J., Trezise, L., Underwood, A., Parsons, M., Kasminkas, L., Rudge, J., Yancopoulos, G., and Vadas, M.A. (2000). Angiopoietin-1 is an antipermeability and anti-inflammatory agent in vitro and targets cell junctions. Circ. Res. 87, 603–607. 10.1161/01.RES.87.7.603.

7. Fiedler, U., and Augustin, H.G. (2006). Angiopoietins: a link between angiogenesis and inflammation. Trends Immunol. 27, 552–558. 10.1016/J.IT.2006.10.004.

8. London, N.R., Whitehead, K.J., and Li, D.Y. (2009). Endogenous endothelial cell signaling systems maintain vascular stability. Angiogenesis 12, 149–158. 10.1007/S10456-009-9130-Z.

9. Huang, H., Bhat, A., Woodnutt, G., and Lappe, R. (2010). Targeting the ANGPT-TIE2 pathway in malignancy. Nat. Rev. Cancer 10, 575–585. 10.1038/NRC2894.

10. Saharinen, P., Leppänen, V.M., and Alitalo, K. (2017). SnapShot: Angiopoietins and Their Functions. Cell 171, 724–724.e1. 10.1016/J.CELL.2017.10.009.

11. Davis, S., Aldrich, T.H., Jones, P.F., Acheson, A., Compton, D.L., Jain, V., Ryan, T.E., Bruno, J., Radziejewski, C., Maisonpierre, P.C., et al. (1996). Isolation of angiopoietin-1, a ligand for the TIE2 receptor, by secretion-trap expression cloning. Cell 87, 1161–1169. 10.1016/S0092-8674(00)81812-7.

12. Suri, C., Jones, P.F., Patan, S., Bartunkova, S., Maisonpierre, P.C., Davis, S., Sato, T.N., and Yancopoulos, G.D. (1996). Requisite role of angiopoietin-1, a ligand for the TIE2 receptor, during embryonic angiogenesis. Cell 87, 1171–1180. 10.1016/S0092-8674(00)81813-9.

13. Akwii, R.G., and Mikelis, C.M. (2021). Targeting the Angiopoietin/Tie Pathway: Prospects for Treatment of Retinal and Respiratory Disorders. Drugs 81, 1731–1749. 10.1007/S40265-021-01605-Y.

14. Gavard, J., Patel, V., and Gutkind, J.S. (2008). Angiopoietin-1 prevents VEGF-induced endothelial permeability by sequestering Src through mDia. Dev. Cell 14, 25–36. 10.1016/J.DEVCEL.2007.10.019.

15. Maisonpierre, P.C., Suri, C., Jones, P.F., Bartunkova, S., Wiegand, S.J., Radziejewski, C., Compton, D., McClain, J., Aldrich, T.H., Papadopoulos, N., et al. (1997). Angiopoietin-2, a natural antagonist for Tie2 that disrupts in vivo angiogenesis. Science 277, 55–60. 10.1126/SCIENCE.277.5322.55.

16. Saharinen, P., Eklund, L., and Alitalo, K. (2017). Therapeutic targeting of the angiopoietin-TIE pathway. Nat. Rev. Drug Discov. 16, 635–661. 10.1038/NRD.2016.278.

17. Siner, J.M., Bhandari, V., Engle, K.M., Elias, J.A., and Siegel, M.D. (2009). Elevated serum angiopoietin 2 levels are associated with increased mortality in sepsis. Shock 31, 348–353. 10.1097/SHK.0B013E318188BD06.

18. Parikh, S.M., Mammoto, T., Schultz, A., Yuan, H.T., Christiani, D., Karumanchi, S.A., and Sukhatme, V.P. (2006). Excess circulating angiopoietin-2 may contribute to pulmonary vascular leak in sepsis in humans. PLoS Med. 3, 356–370. 10.1371/JOURNAL.PMED.0030046.

19. Agrawal, A., Matthay, M.A., Kangelaris, K.N., Stein, J., Chu, J.C., Imp, B.M., Cortez, A., Abbott, J., Liu, K.D., and Calfee, C.S. (2013). Plasma angiopoietin-2 predicts the onset of acute lung injury in critically ill patients. Am. J. Respir. Crit. Care Med. 187, 736–742. 10.1164/RCCM.201208-1460OC.

20. Parikh, S.M. (2013). Dysregulation of the angiopoietin-Tie-2 axis in sepsis and ARDS. Virulence 4. 10.4161/VIRU.24906.

21. Cho, C.H., Kammerer, R.A., Lee, H.J., Steinmetz, M.O., Ryu, Y.S., Lee, S.H., Yasunaga, K., Kim, K.T., Kim, I., Choi, H.H., et al. (2004). COMP-Ang1: a designed angiopoietin-1 variant with nonleaky angiogenic activity. Proc. Natl. Acad. Sci. U. S. A. 101, 5547– 5552. 10.1073/PNAS.0307574101.

22. Oh, N., Kim, K., Jin Kim, S., Park, I., Lee, J.E., Suk Seo, Y., Joo An, H., Min Kim, H., and Young Koh, G. (2015). A Designed Angiopoietin-1 Variant, Dimeric CMP-Ang1 Activates Tie2 and Stimulates Angiogenesis and Vascular Stabilization in N-glycan Dependent Manner. Sci. Rep. 5. 10.1038/SREP15291.

23. Alfieri, A., Watson, J.J., Kammerer, R.A., Tasab, M., Progias, P., Reeves, K., Brown, N.J., and Brookes, Z.L. (2012). Angiopoietin-1 variant reduces LPS-induced microvascular dysfunction in a murine model of sepsis. Crit. Care 16. 10.1186/CC11666.

24. Wallace, R.G., Rochfort, K.D., Barabas, P., Curtis, T.M., Uehara, H., Ambati, B.K., and Cummins, P.M. (2021). COMP-Ang1: Therapeutic potential of an engineered Angiopoietin-1 variant. Vascul. Pharmacol. 141. 10.1016/J.VPH.2021.106919.

25. Liu, P., Ryczko, M., Xie, X., Baardsnes, J., Lord-Dufour, S., Duroche, Y., Hicks, E.A., Taiyab, A., Sheardown, H., Quaggin, S.E., et al. (2021). New soluble angiopoietin analog of Hepta-ANG1 prevents pathological vascular leakage. Biotechnol. Bioeng. 118, 423–432. 10.1002/BIT.27580.

26. Liu, F.Y., Cho, Y.L., Fridayana, F.R., Niloofar, L., Vo, M.N., Huang, Y., Limanjaya, A., Kwon, M.H., Ock, J., Lee, S.J., et al. (2025). MT-100, a human Tie2-agonistic antibody, improves penile neurovasculature in diabetic mice via the novel target Srpx2. Exp. Mol. Med. 57, 104–117. 10.1038/S12276-024-01373-1.

27. Jo, G., Bae, J., Hong, H.J., Han, A. reum, Kim, D.K., Hong, S.P., Kim, J.A., Lee, S., Koh, G.Y., and Kim, H.M. (2021). Structural insights into the clustering and activation of Tie2 receptor mediated by Tie2 agonistic antibody. Nat. Commun. 12. 10.1038/S41467-021-26620-1.

28. Van Slyke, P., Alami, J., Martin, D., Kuliszewski, M., Leong-Poi, H., Sefton, M. V., and Dumont, D. (2009). Acceleration of diabetic wound healing by an angiopoietin peptide mimetic. Tissue Eng. Part A 15, 1269–1280. 10.1089/TEN.TEA.2007.0400.

29. Kümpers, P., Gueler, F., David, S., Van Slyke, P., Dumont, D.J., Park, J.K., Bockmeyer, C.L., Parikh, S.M., Pavenstädt, H., Haller, H., et al. (2011). The synthetic Tie2 agonist peptide vasculotide protects against vascular leakage and reduces mortality in murine abdominal sepsis. Crit. Care 15. 10.1186/cc10523.

30. Lomas-Neira, J.L., Heffernan, D.S., Ayala, A., and Monaghan, S.F. (2016). BLOCKADE OF ENDOTHELIAL GROWTH FACTOR, ANGIOPOIETIN-2, REDUCES INDICES OF ARDS AND MORTALITY IN MICE RESULTING FROM THE DUAL-INSULTS OF HEMORRHAGIC SHOCK AND SEPSIS. Shock 45, 157–165. 10.1097/SHK.0000000000000499.

31. David, S., Mukherjee, A., Ghosh, C.C., Yano, M., Khankin, E. V., Wenger, J.B., Ananth Karumanchi, S., Shapiro, N.I., and Parikh, S.M. (2012). Angiopoietin-2 may contribute to multiple organ dysfunction and death in sepsis*. Crit. Care Med. 40, 3034–3041. 10.1097/CCM.0B013E31825FDC31.

32. Han, S., Lee, S.J., Kim, K.E., Lee, H.S., Oh, N., Park, I., Ko, E., Oh, S.J., Lee, Y.S., Kim, D., et al. (2016). Amelioration of sepsis by TIE2 activation-induced vascular protection. Sci. Transl. Med. 8. 10.1126/SCITRANSLMED.AAD9260.

33. Maisonpierre, P.C., Suri, C., Jones, P.F., Bartunkova, S., Wiegand, S.J., Radziejewski, C., Compton, D., McClain, J., Aldrich, T.H., Papadopoulos, N., et al. (1997). Angiopoietin-2, a natural antagonist for Tie2 that disrupts in vivo angiogenesis. Science 277, 55–60. 10.1126/SCIENCE.277.5322.55.

34. Akwii, R.G., Sajib, S., Zahra, F.T., and Mikelis, C.M. (2019). Role of Angiopoietin-2 in Vascular Physiology and Pathophysiology. Cells 8. 10.3390/CELLS8050471.

35. Reusch, P., Barleon, B., Weindel, K., Martiny-Baron, G., Gödde, A., Siemeister, G., and Marmé, D. (2001). Identification of a soluble form of the angiopoietin receptor TIE-2 released from endothelial cells and present in human blood. Angiogenesis 4, 123–131. 10.1023/A:1012226627813.

36. Findley, C.M., Cudmore, M.J., Ahmed, A., and Kontos, C.D. (2007). VEGF induces Tie2 shedding via a phosphoinositide 3-kinase/Akt dependent pathway to modulate Tie2 signaling. Arterioscler. Thromb. Vasc. Biol. 27, 2619–2626. 10.1161/ATVBAHA.107.150482.

37. Van Der Heijden, M., Van Nieuw Amerongen, G.P., Van Hinsbergh, V.W.M., and Groeneveld, A.J. (2010). The interaction of soluble Tie2 with angiopoietins and pulmonary vascular permeability in septic and nonseptic critically ill patients. Shock 33, 263–268. 10.1097/SHK.0B013E3181B2F978.

38. Idowu, T.O., Etzrodt, V., Seeliger, B., Bolanos-Palmieri, P., Thamm, K., Haller, H., and David, S. (2020). Identification of specific Tie2 cleavage sites and therapeutic modulation in experimental sepsis. Elife 9, 1–15. 10.7554/ELIFE.59520.

39. Fachinger, G., Deutsch, U., and Risau, W. (1999). Functional interaction of vascular endothelial-protein-tyrosine phosphatase with the angiopoietin receptor Tie-2. Oncogene 18, 5948–5953. 10.1038/SJ.ONC.1202992.

40. Winderlich, M., Keller, L., Cagna, G., Broermann, A., Kamenyeva, O., Kiefer, F., Deutsch, U., Nottebaum, A.F., and Vestweber, D. (2009). VE-PTP controls blood vessel development by balancing Tie-2 activity. Journal of Cell Biology 185, 657–671. 10.1083/jcb.200811159.

41. Schmaier, A.A., Pajares Hurtado, G.M., Manickas-Hill, Z.J., Sack, K.D., Chen, S.M., Bhambhani, V., Quadir, J., Nath, A.K., Collier, A.R.Y., Ngo, D., et al. (2021). Tie2 activation protects against prothrombotic endothelial dysfunction in COVID-19. JCI Insight 6. 10.1172/JCI.INSIGHT.151527.

42. Study Details | NCT02050828 | The TIME-2 Study: A Phase 2 Study of AKB-9778, a Novel Tie-2 Activator, in Patients With Diabetic Macular Edema | ClinicalTrials.gov https://clinicaltrials.gov/study/NCT02050828.

43. Wehrle, C., Van Slyke, P., and Dumont, D.J. (2009). Angiopoietin-1-induced ubiquitylation of Tie2 by c-Cbl is required for internalization and degradation. Biochem. J. 423, 375–380. 10.1042/BJ20091010.

44. Radloff, K., Gutbier, B., Dunne, C.M., Moradian, H., Schwestka, M., Gossen, M., Ahrens, K., Kneller, L., Wang, Y., Moga, A., et al. (2023). Cationic LNP-formulated mRNA expressing Tie2-agonist in the lung endothelium prevents pulmonary vascular leakage. Mol. Ther. Nucleic Acids 34. 10.1016/J.OMTN.2023.102068.

45. Critchley, W.R., Pellet-Many, C., Ringham-Terry, B., Harrison, M.A., Zachary, I.C., and Ponnambalam, S. (2018). Receptor Tyrosine Kinase Ubiquitination and De-Ubiquitination in Signal Transduction and Receptor Trafficking. Cells 7. 10.3390/CELLS7030022.

46. Mohapatra, B., Ahmad, G., Nadeau, S., Zutshi, N., An, W., Scheffe, S., Dong, L., Feng, D., Goetz, B., Arya, P., et al. (2013). Protein tyrosine kinase regulation by ubiquitination: critical roles of Cbl-family ubiquitin ligases. Biochim. Biophys. Acta 1833, 122–139. 10.1016/J.BBAMCR.2012.10.010.

47. Hu, H.T., Huang, Y.H., Chang, Y.A., Lee, C.K., Jiang, M.J., and Wu, L.W. (2008). Tie2-R849W mutant in venous malformations chronically activates a functional STAT1 to modulate gene expression. J. Invest. Dermatol. 128, 2325–2333. 10.1038/JID.2008.89.

48. Kennedy, M.A., Xu, Z., Wu, Y., and Sohl, C.D. (2019). A Tie2 kinase mutation causing venous malformations increases phosphorylation rates and enhances cooperativity. Biochem. Biophys. Res. Commun. 509, 898–902. 10.1016/J.BBRC.2019.01.020.

49. Soblet, J., Limaye, N., Uebelhoer, M., Boon, L.M., and Vikkula, M. (2013). Variable Somatic TIE2 Mutations in Half of Sporadic Venous Malformations. Mol. Syndromol. 4, 179–183. 10.1159/000348327.

50. Limaye, N., Wouters, V., Uebelhoer, M., Tuominen, M., Wirkkala, R., Mulliken, J.B., Eklund, L., Boon, L.M., and Vikkula, M. (2009). Somatic mutations in angiopoietin receptor gene TEK cause solitary and multiple sporadic venous malformations. Nat. Genet. 41, 118–124. 10.1038/NG.272.

51. Uebelhoer, M., Nätynki, M., Kangas, J., Mendola, A., Nguyen, H.L., Soblet, J., Godfraind, C., Boon, L.M., Eklund, L., Limaye, N., et al. (2013). Venous malformation-causative TIE2 mutations mediate an AKT-dependent decrease in PDGFB. Hum. Mol. Genet. 22, 3438–3448. 10.1093/HMG/DDT198.

52. Potente, M., Urbich, C., Sasaki, K.I., Hofmann, W.K., Heeschen, C., Aicher, A., Kollipara, R., DePinho, R.A., Zeiher, A.M., and Dimmeler, S. (2005). Involvement of Foxo transcription factors in angiogenesis and postnatal neovascularization. J. Clin. Invest. 115, 2382–2392. 10.1172/JCI23126.

53. Cain, R.J., Vanhaesebroeck, B., and Ridley, A.J. (2010). The PI3K p110alpha isoform regulates endothelial adherens junctions via Pyk2 and Rac1. J. Cell Biol. 188, 863–876. 10.1083/JCB.200907135.

54. Huveneers, S., Oldenburg, J., Spanjaard, E., van der Krogt, G., Grigoriev, I., Akhmanova, A., Rehmann, H., and de Rooij, J. (2012). Vinculin associates with endothelial VE-cadherin junctions to control force-dependent remodeling. J. Cell Biol. 196, 641–652. 10.1083/JCB.201108120.

55. Angelidis, I., Simon, L.M., Fernandez, I.E., Strunz, M., Mayr, C.H., Greiffo, F.R., Tsitsiridis, G., Ansari, M., Graf, E., Strom, T.M., et al. (2019). An atlas of the aging lung mapped by single cell transcriptomics and deep tissue proteomics. Nat. Commun. 10. 10.1038/S41467-019-08831-9.

56. Gillich, A., Zhang, F., Farmer, C.G., Travaglini, K.J., Tan, S.Y., Gu, M., Zhou, B., Feinstein, J.A., Krasnow, M.A., and Metzger, R.J. (2020). Capillary cell-type specialization in the alveolus. Nature 586, 785–789. 10.1038/S41586-020-2822-7.

57. Donati, Y., Blaskovic, S., Ruchonnet-Métrailler, I., Maillard, J.L., and Barazzone-Argiroffo, C. (2020). Simultaneous isolation of endothelial and alveolar epithelial type I and type II cells during mouse lung development in the absence of a transgenic reporter. Am. J. Physiol. Lung Cell. Mol. Physiol. 318, L619–L630. 10.1152/AJPLUNG.00227.2019.

58. Parikh, S.M. (2017). Angiopoietins and Tie2 in vascular inflammation. Curr. Opin. Hematol. 24, 432–438. 10.1097/MOH.0000000000000361.

59. Yuan, H.T., Khankin, E. V., Karumanchi, S.A., and Parikh, S.M. (2009). Angiopoietin 2 is a partial agonist/antagonist of Tie2 signaling in the endothelium. Mol. Cell. Biol. 29, 2011–2022. 10.1128/MCB.01472-08.

60. Sack, K.D., Kellum, J.A., and Parikh, S.M. (2020). The Angiopoietin-Tie2 Pathway in Critical Illness. Crit. Care Clin. 36, 201–216. 10.1016/J.CCC.2019.12.003.

61. Latreille, E., and Lee, W.L. (2021). Interactions of Influenza and SARS-CoV-2 with the Lung Endothelium: Similarities, Differences, and Implications for Therapy. Viruses 13. 10.3390/V13020161.

62. Smadja, D.M., Guerin, C.L., Chocron, R., Yatim, N., Boussier, J., Gendron, N., Khider, L., Hadjadj, J., Goudot, G., Debuc, B., et al. (2020). Angiopoietin-2 as a marker of endothelial activation is a good predictor factor for intensive care unit admission of COVID-19 patients. Angiogenesis 23, 611–620. 10.1007/S10456-020-09730-0.

63. Koenecke, C., Kümpers, P., Lukasz, A., Dammann, E., Verhagen, W., Göhring, G., Buchholz, S., Krauter, J., Eder, M., Schlegelberger, B., et al. (2010). Shedding of the endothelial receptor tyrosine kinase Tie2 correlates with leukemic blast burden and outcome after allogeneic hematopoietic stem cell transplantation for AML. Ann. Hematol. 89, 459–467. 10.1007/S00277-009-0869-5.

64. Wegmann, N.J., Heuberger, D.M., Kadler, G., Seeliger, B., Pape, T., Stahl, K., Müller, M.M., and David, S. (2025). Association of the soluble form of the endothelial Tie2 receptor with organ failure and mortality in patients with sepsis and ARDS. Cytokine 196. 10.1016/J.CYTO.2025.157062.

65. Jo, G., Bae, J., Hong, H.J., Han, A. reum, Kim, D.K., Hong, S.P., Kim, J.A., Lee, S., Koh, G.Y., and Kim, H.M. (2021). Structural insights into the clustering and activation of Tie2 receptor mediated by Tie2 agonistic antibody. Nat. Commun. 12. 10.1038/S41467-021-26620-1.

66. Han, S., Lee, S.J., Kim, K.E., Lee, H.S., Oh, N., Park, I., Ko, E., Oh, S.J., Lee, Y.S., Kim, D., et al. (2016). Amelioration of sepsis by TIE2 activation-induced vascular protection. Sci. Transl. Med. 8. 10.1126/SCITRANSLMED.AAD9260.

67. Pietilä, R., Marks-Hultström, A., He, L., Nanavazadeh, S., Quaggin, S.E., Betsholtz, C., and Jeansson, M. (2025). TIE2 activation by antibody-clustered endogenous angiopoietin-2 prevents capillary loss and fibrosis in experimental kidney disease. J. Clin. Invest. 135. 10.1172/JCI190286.

68. Saharinen, P., Eklund, L., Miettinen, J., Wirkkala, R., Anisimov, A., Winderlich, M., Nottebaum, A., Vestweber, D., Deutsch, U., Koh, G.Y., et al. (2008). Angiopoietins assemble distinct Tie2 signalling complexes in endothelial cell-cell and cell-matrix contacts. Nat. Cell Biol. 10, 527–537. 10.1038/NCB1715.

69. Fukuhara, S., Sako, K., Minami, T., Noda, K., Kim, H.Z., Kodama, T., Shibuya, M., Takakura, N., Koh, G.Y., and Mochizuki, N. (2008). Differential function of Tie2 at cell-cell contacts and cell-substratum contacts regulated by angiopoietin-1. Nat. Cell Biol. 10, 513–526. 10.1038/NCB1714.

70. Fukuhara, S., Sako, K., Noda, K., Nagao, K., Miura, K., and Mochizuki, N. (2009). Tie2 is tied at the cell-cell contacts and to extracellular matrix by angiopoietin-1. Exp. Mol. Med. 41, 133–139. 10.3858/EMM.2009.41.3.016.

71. Thomas, M., Felcht, M., Kruse, K., Kretschmer, S., Deppermann, C., Biesdorf, A., Rohr, K., Benest, A. V., Fiedler, U., and Augustin, H.G. (2010). Angiopoietin-2 stimulation of endothelial cells induces alphavbeta3 integrin internalization and degradation. J. Biol. Chem. 285, 23842–23849. 10.1074/JBC.M109.097543.

72. Hakanpaa, L., Sipila, T., Leppanen, V.M., Gautam, P., Nurmi, H., Jacquemet, G., Eklund, L., Ivaska, J., Alitalo, K., and Saharinen, P. (2015). Endothelial destabilization by angiopoietin-2 via integrin β1 activation. Nat. Commun. 6. 10.1038/NCOMMS6962.

73. Fukuhara, S., Sako, K., Noda, K., Nagao, K., Miura, K., and Mochizuki, N. (2009). Tie2 is tied at the cell-cell contacts and to extracellular matrix by angiopoietin-1. Exp. Mol. Med. 41, 133–139. 10.3858/EMM.2009.41.3.016.

74. Klippel, A., Reinhard, C., Kavanaugh, W.M., Apell, G., Escobedo, M.-A., and Williams, L.T. (1996). Membrane localization of phosphatidylinositol 3-kinase is sufficient to activate multiple signal-transducing kinase pathways. Mol. Cell. Biol. 16, 4117–4127. 10.1128/MCB.16.8.4117.

75. Baumgartner, W., Weth, A., Gutberlet, J., Harms, G., and Groschner, K. (2014). Localization of VE-cadherin in plasmalemmal cholesterol rich microdomains and the effects of cholesterol depletion on VE-cadherin mediated cell-cell adhesion. Biochim. Biophys. Acta 1841, 1725–1732. 10.1016/J.BBALIP.2014.08.016.

76. Irwin, M.E., Bohin, N., and Boerner, J.L. (2011). Src family kinases mediate epidermal growth factor receptor signaling from lipid rafts in breast cancer cells. Cancer Biol. Ther. 12, 718–726. 10.4161/CBT.12.8.16907.

77. Webster, M.K., and Donoghue, D.J. (1996). Constitutive activation of fibroblast growth factor receptor 3 by the transmembrane domain point mutation found in achondroplasia. EMBO J. 15, 520. 10.1002/j.1460-2075.1996.tb00384.x.

78. Hart, K.C., Robertson, S.C., and Donoghue, D.J. (2001). Identification of Tyrosine Residues in Constitutively Activated Fibroblast Growth Factor Receptor 3 Involved in Mitogenesis, Stat Activation, and Phosphatidylinositol 3-Kinase Activation. Mol. Biol. Cell 12, 931. 10.1091/MBC.12.4.931.

79. Terskikh, A. V., Le Doussal, J.M., Crameri, R., Fisch, I., Mach, J.P., and Kajava, A. V. (1997). “Peptabody”: a new type of high avidity binding protein. Proc. Natl. Acad. Sci. U. S. A. 94, 1663–1668. 10.1073/PNAS.94.5.1663.

80. Kim, I., Kim, H.G., So, J.N., Kim, J.H., Kwak, H.J., and Koh, G.Y. (2000). Angiopoietin-1 regulates endothelial cell survival through the phosphatidylinositol 3’-kinase/Akt signal transduction pathway. Circ. Res. 86, 24–29. 10.1161/01.RES.86.1.24.

81. Daly, C., Wong, V., Burova, E., Wei, Y., Zabski, S., Griffiths, J., Lai, K.M., Lin, H.C., Ioffe, E., Yancopoulos, G.D., et al. (2004). Angiopoietin-1 modulates endothelial cell function and gene expression via the transcription factor FKHR (FOXO1). Genes Dev. 18, 1060–1071. 10.1101/GAD.1189704.

82. Potente, M., Urbich, C., Sasaki, K.I., Hofmann, W.K., Heeschen, C., Aicher, A., Kollipara, R., DePinho, R.A., Zeiher, A.M., and Dimmeler, S. (2005). Involvement of Foxo transcription factors in angiogenesis and postnatal neovascularization. J. Clin. Invest. 115, 2382–2392. 10.1172/JCI23126.

83. Kim, K.H., Nakaoka, Y., Augustin, H.G., and Koh, G.Y. (2018). Myocardial Angiopoietin-1 Controls Atrial Chamber Morphogenesis by Spatiotemporal Degradation of Cardiac Jelly. Cell Rep. 23, 2455–2466. 10.1016/J.CELREP.2018.04.080.

84. Nätynki, M., Kangas, J., Miinalainen, I., Sormunen, R., Pietilä, R., Soblet, J., Boon, L.M., Vikkula, M., Limaye, N., and Eklund, L. (2015). Common and specific effects of TIE2 mutations causing venous malformations. Hum. Mol. Genet. 24, 6374–6389. 10.1093/HMG/DDV349.

85. Si, Y., Huang, J., Li, X., Fu, Y., Xu, R., Du, Y., Cheng, J., and Jiang, H. (2020). AKT/FOXO1 axis links cross-talking of endothelial cell and pericyte in TIE2-mutated venous malformations. Cell Commun. Signal. 18. 10.1186/S12964-020-00606-W.

86. Uebelhoer, M., Nätynki, M., Kangas, J., Mendola, A., Nguyen, H.L., Soblet, J., Godfraind, C., Boon, L.M., Eklund, L., Limaye, N., et al. (2013). Venous malformation-causative TIE2 mutations mediate an AKT-dependent decrease in PDGFB. Hum. Mol. Genet. 22, 3438–3448. 10.1093/HMG/DDT198.

87. Sako, K., Fukuhara, S., Minami, T., Hamakubo, T., Song, H., Kodama, T., Fukamizu, A., Gutkind, J.S., Koh, G.Y., and Mochizuki, N. (2009). Angiopoietin-1 induces Kruppel-like factor 2 expression through a phosphoinositide 3-kinase/AKT-dependent activation of myocyte enhancer factor 2. J. Biol. Chem. 284, 5592–5601. 10.1074/JBC.M806928200.

88. Zhang, J., Fukuhara, S., Sako, K., Takenouchi, T., Kitani, H., Kume, T., Koh, G.Y., and Mochizuki, N. (2011). Angiopoietin-1/Tie2 signal augments basal Notch signal controlling vascular quiescence by inducing delta-like 4 expression through AKT-mediated activation of beta-catenin. J. Biol. Chem. 286, 8055–8066. 10.1074/JBC.M110.192641.

89. Yan, M., and Plowman, G.D. (2007). Delta-like 4/Notch signaling and its therapeutic implications. Clin. Cancer Res. 13, 7243–7246. 10.1158/1078-0432.CCR-07-1393.

90. Kume, T. (2012). Ligand-dependent Notch signaling in vascular formation. Adv. Exp. Med. Biol. 727, 210–222. 10.1007/978-1-4614-0899-4_16.

91. Benedito, R., Roca, C., Sörensen, I., Adams, S., Gossler, A., Fruttiger, M., and Adams, R.H. (2009). The Notch Ligands Dll4 and Jagged1 Have Opposing Effects on Angiogenesis. Cell 137, 1124–1135. 10.1016/j.cell.2009.03.025.

92. Blanco, R., and Gerhardt, H. (2013). VEGF and Notch in tip and stalk cell selection. Cold Spring Harb. Perspect. Med. 3. 10.1101/CSHPERSPECT.A006569.

93. Iruela-Arispe, M.L., Carpizo, D., and Luque, A. (2003). ADAMTS1: a matrix metalloprotease with angioinhibitory properties. Ann. N. Y. Acad. Sci. 995, 183–190. 10.1111/J.1749-6632.2003.TB03221.X.

94. Luque, A., Carpizo, D.R., and Iruela-Arispe, M.L. (2003). ADAMTS1/METH1 inhibits endothelial cell proliferation by direct binding and sequestration of VEGF165. J. Biol. Chem. 278, 23656–23665. 10.1074/JBC.M212964200.

95. Vázquez, F., Hastings, G., Ortega, M.A., Lane, T.F., Oikemus, S., Lombardo, M., and Iruela-Arispe, M.L. (1999). METH-1, a human ortholog of ADAMTS-1, and METH-2 are members of a new family of proteins with angio-inhibitory activity. J. Biol. Chem. 274, 23349–23357. 10.1074/JBC.274.33.23349.

96. Daly, C., Wong, V., Burova, E., Wei, Y., Zabski, S., Griffiths, J., Lai, K.M., Lin, H.C., Ioffe, E., Yancopoulos, G.D., et al. (2004). Angiopoietin-1 modulates endothelial cell function and gene expression via the transcription factor FKHR (FOXO1). Genes Dev. 18, 1060–1071. 10.1101/GAD.1189704.

97. Fukuhara, S., Sako, K., Minami, T., Noda, K., Kim, H.Z., Kodama, T., Shibuya, M., Takakura, N., Koh, G.Y., and Mochizuki, N. (2008). Differential function of Tie2 at cell-cell contacts and cell-substratum contacts regulated by angiopoietin-1. Nat. Cell Biol. 10, 513–526. 10.1038/NCB1714.

98. Uebelhoer, M., Nätynki, M., Kangas, J., Mendola, A., Nguyen, H.L., Soblet, J., Godfraind, C., Boon, L.M., Eklund, L., Limaye, N., et al. (2013). Venous malformation-causative TIE2 mutations mediate an AKT-dependent decrease in PDGFB. Hum. Mol. Genet. 22, 3438–3448. 10.1093/HMG/DDT198.

99. Whyte, C.S., Morrow, G.B., Mitchell, J.L., Chowdary, P., and Mutch, N.J. (2020). Fibrinolytic abnormalities in acute respiratory distress syndrome (ARDS) and versatility of thrombolytic drugs to treat COVID-19. J. Thromb. Haemost. 18, 1548–1555. 10.1111/JTH.14872.

100. C. Sebag, S., A. Bastarache, J., and B. Ware, L. (2011). Therapeutic modulation of coagulation and fibrinolysis in acute lung injury and the acute respiratory distress syndrome. Curr. Pharm. Biotechnol. 12, 1481–1496. 10.2174/138920111798281171.

101. Isabel García-Laorden, M., Lorente, J.A., Flores, C., Slutsky, A.S., and Villar, J. (2017). Biomarkers for the acute respiratory distress syndrome: how to make the diagnosis more precise. Ann. Transl. Med. 5. 10.21037/ATM.2017.06.49.

102. Van Der Zee, P., Rietdijk, W., Somhorst, P., Endeman, H., and Gommers, D. (2020). A systematic review of biomarkers multivariately associated with acute respiratory distress syndrome development and mortality. Crit. Care 24. 10.1186/S13054-020-02913-7.

103. Khatri, S., Suchday, P., Guddeti, A., Nanna, S., Gupta, S., Darapaneni, H., Mohammed, A.S., Desai, R., and Imtiaz, H. (2025). Endocan as a Novel Biomarker for Endothelial Dysfunction and Cardiovascular Prognosis in ST-Elevation Myocardial Infarction: A Contemporary Literature Review. J. Pers. Med. 16, 7. 10.3390/JPM16010007.

104. Bidlan, M., Kumar, M., Bhargava, A., Goel, V., and Kumar, V. (2025). Endocan, a Potential Biomarker of Endothelial Dysfunction: A Brief Overview. Angiology. 10.1177/00033197251356576.

105. Kular, L., Pakradouni, J., Kitabgi, P., Laurent, M., and Martinerie, C. (2011). The CCN family: a new class of inflammation modulators? Biochimie 93, 377–388. 10.1016/J.BIOCHI.2010.11.010.

106. Kubota, S., and Takigawa, M. (2007). CCN family proteins and angiogenesis: from embryo to adulthood. Angiogenesis 10, 1–11. 10.1007/S10456-006-9058-5.

107. Babic, A.M., Kireeva, M.L., Kolesnikova, T. V., and Lau, L.F. (1998). CYR61, a product of a growth factor-inducible immediate early gene, promotes angiogenesis and tumor growth. Proc. Natl. Acad. Sci. U. S. A. 95, 6355–6360. 10.1073/PNAS.95.11.6355.

108. Hall-Glenn, F., de Young, R.A., Huang, B.L., van Handel, B., Hofmann, J.J., Chen, T.T., Choi, A., Ong, J.R., Benya, P.D., Mikkola, H., et al. (2012). CCN2/connective tissue growth factor is essential for pericyte adhesion and endothelial basement membrane formation during angiogenesis. PLoS One 7. 10.1371/JOURNAL.PONE.0030562.

109. Wilhelm, K., Happel, K., Eelen, G., Schoors, S., Oellerich, M.F., Lim, R., Zimmermann, B., Aspalter, I.M., Franco, C.A., Boettger, T., et al. (2016). FOXO1 couples metabolic activity and growth state in the vascular endothelium. Nature 529, 216–220. 10.1038/NATURE16498.

110. Dharaneeswaran, H., Abid, M.R., Yuan, L., Dupuis, D., Beeler, D., Spokes, K.C., Janes, L., Sciuto, T., Kang, P.M., Jaminet, S.C.S., et al. (2014). FOXO1-mediated activation of Akt plays a critical role in vascular homeostasis. Circ. Res. 115, 238–251. 10.1161/CIRCRESAHA.115.303227.

111. Li-Villarreal, N., Wong, R.L.Y., Garcia, M.D., Udan, R.S., Poché, R.A., Rasmussen, T.L., Rhyner, A.M., Wythe, J.D., and Dickinson, M.E. (2022). FOXO1 represses sprouty 2 and sprouty 4 expression to promote arterial specification and vascular remodeling in the mouse yolk sac. Development 149, dev200131. 10.1242/DEV.200131.

112. Suchting, S., Freitas, C., Le Noble, F., Benedito, R., Bréant, C., Duarte, A., and Eichmann, A. (2007). The Notch ligand Delta-like 4 negatively regulates endothelial tip cell formation and vessel branching. Proc. Natl. Acad. Sci. U. S. A. 104, 3225–3230. 10.1073/PNAS.0611177104/SUPPL_FILE/11177FIG7.JPG.

113. Williams, C.K., Li, J.L., Murga, M., Harris, A.L., and Tosato, G. (2005). Up-regulation of the Notch ligand Delta-like 4 inhibits VEGF-induced endothelial cell function. Blood 107, 931. 10.1182/BLOOD-2005-03-1000.

114. Pitulescu, M.E., Schmidt, I., Giaimo, B.D., Antoine, T., Berkenfeld, F., Ferrante, F., Park, H., Ehling, M., Biljes, D., Rocha, S.F., et al. (2017). Dll4 and Notch signalling couples sprouting angiogenesis and artery formation. Nat. Cell Biol. 19, 915–927. 10.1038/NCB3555.

115. Potente, M., Urbich, C., Sasaki, K.I., Hofmann, W.K., Heeschen, C., Aicher, A., Kollipara, R., DePinho, R.A., Zeiher, A.M., and Dimmeler, S. (2005). Involvement of Foxo transcription factors in angiogenesis and postnatal neovascularization. J. Clin. Invest. 115, 2382–2392. 10.1172/JCI23126.

116. Thines, L., Roushar, F.J., Hedman, A.C., and Sacks, D.B. (2023). The IQGAP scaffolds: Critical nodes bridging receptor activation to cellular signaling. J. Cell Biol. 222. 10.1083/JCB.202205062.

117. David, S., Ghosh, C.C., Mukherjee, A., and Parikh, S.M. (2011). Angiopoietin-1 requires IQ domain GTPase-activating protein 1 to activate Rac1 and promote endothelial barrier defense. Arterioscler. Thromb. Vasc. Biol. 31, 2643–2652. 10.1161/ATVBAHA.111.233189.

118. Yamaoka-Tojo, M., Tojo, T., Kim, H.W., Hilenski, L., Patrushev, N.A., Zhang, L., Fukai, T., and Ushio-Fukai, M. (2006). IQGAP1 mediates VE-cadherin-based cell-cell contacts and VEGF signaling at adherence junctions linked to angiogenesis. Arterioscler. Thromb. Vasc. Biol. 26, 1991–1997. 10.1161/01.ATV.0000231524.14873.E7/SUPPL_FILE/ATVB7455-SUPPL.PDF.

119. Yamaoka-Tojo, M., Ushio-Fukai, M., Hilenski, L., Dikalov, S.I., Chen, Y.E., Tojo, T., Fukai, T., Fujimoto, M., Patrushev, N.A., Wang, N., et al. (2004). IQGAP1, a novel vascular endothelial growth factor receptor binding protein, is involved in reactive oxygen species--dependent endothelial migration and proliferation. Circ. Res. 95, 276–283. 10.1161/01.RES.0000136522.58649.60.

120. Thickett, D.R., Armstrong, L., Christie, S.J., and Millar, A.B. (2001). Vascular endothelial growth factor may contribute to increased vascular permeability in acute respiratory distress syndrome. Am. J. Respir. Crit. Care Med. 164, 1601–1605. 10.1164/AJRCCM.164.9.2011071.

